# Deep Generative Optimization of mRNA Codon Sequences for Enhanced mRNA Translation and Therapeutic Efficacy

**DOI:** 10.1101/2024.09.06.611590

**Authors:** Yupeng Li, Fan Wang, Jiaqi Yang, Zirong Han, Linfeng Chen, Wenbing Jiang, Hao Zhou, Tong Li, Zehua Tang, Jianxiang Deng, Xin He, Gaofeng Zha, Jiekai Hu, Yong Hu, Linping Wu, Changyou Zhan, Caijun Sun, Yao He, Zhi Xie

## Abstract

Messenger RNA (mRNA) therapeutics show immense promise, but their efficacy is limited by suboptimal protein expression. Here, we present RiboDecode, a deep learning framework that generates mRNA codon sequences for enhanced mRNA translation. RiboDecode introduces several advances, including direct learning from large-scale ribosome profiling data and generative exploration of a large sequence space. *In silico* analysis demonstrates RiboDecode’ s robust predictive accuracy for unseen genes and cellular environments. *In vitro* experiments showed substantial improvements in protein expression, significantly outperforming past methods. In addition, RiboDecode enables mRNA design with consideration of cellular context and demonstrates robust performance across different mRNA formats, including m^1^Ψ-modified and circular mRNAs, an important feature for mRNA therapeutics. *In vivo* mouse studies showed that optimized influenza hemagglutinin mRNAs induce ten times stronger neutralizing antibody responses against influenza virus compared to the unoptimized sequence. In an optic nerve crush model, optimized nerve growth factor mRNAs achieve equivalent neuroprotection of retinal ganglion cells at one-fifth the dose of the unoptimized sequence. Collectively, RiboDecode represents a paradigm shift from rule-based to a data-driven, context-aware approach for mRNA therapeutic applications, enabling the development of more potent and dose-efficient treatments.

## INTRODUCTION

Messenger RNA (mRNA) therapy has emerged as a promising approach for treating diseases. This innovative therapeutic strategy harnesses the cell’s protein synthesis machinery to produce desired proteins encoded by the delivered mRNA^1–3^, leading to the application of mRNA therapies in various fields, such as vaccine development and protein replacement therapy^4^. The successful development and deployment of mRNA vaccines during the COVID-19 pandemic have further highlighted the transformative potential of this technology^5^.

Despite the remarkable progress in mRNA vaccines, achieving efficient and consistent protein translation from delivered mRNA molecules remains a key challenge, particularly critical for protein replacement therapy where sustained, precise, and often higher levels of protein expression are required in specific cellular contexts. However, the biological instability of mRNA and the complex regulatory mechanisms governing mRNA translation in cells can lead to suboptimal protein expression^6–8^. Therefore, improving the expression of mRNA is a key challenge for enhancing the therapeutic efficacy and reducing the required dose of mRNA-based treatments.

An amino acid can be encoded by multiple synonymous codons, ranging from one to six codons per amino acid. Codon optimization is a strategy to improve protein expression by changing the synonymous codon of an mRNA molecule while maintaining the encoded amino acid sequence. The choice of synonymous codons can largely impact the efficiency of mRNA translation and the stability of the mRNA molecule^6,7^. For example, it has been shown that optimal codon usage can enhance ribosome engagement and increase translation elongation rates, ultimately leading to higher protein production^8^. Additionally, codon choice can influence mRNA structure. Previous studies have demonstrated that mRNA structure critically influences its stability *in vivo*^9^, in solution^10^, and translation^11^. Therefore, codon optimization is a critical step in the design of mRNA-based therapies to achieve maximal protein production, leading to better therapeutic efficacy.

Computational tools have been developed for codon optimization, most of which were designed for DNA, employing various strategies to select optimal codons. Past methods rely on codon usage bias derived from highly expressed genes in a given species, such as codon adaptation index (CAI)^12^. These methods aim to mimic the codon usage patterns of efficiently translated endogenous mRNAs. More recently, LinearDesign has been developed for mRNA optimization, aiming to jointly optimize translation and mRNA stability by increasing CAI and reducing minimum free energy (MFE)^6^, which is a computational metric for evaluating mRNA secondary structure. LinearDesign uses a linear programming approach to explore a wider space of sequence variants compared to previous methods and showed superior performance over the previous codon optimization methods. Additionally, other indices have been used to guide sequence optimization. For instance, higher GC content (GC%) has been associated with enhanced gene expression^13^.

Despite the development of the previous methods, several limitations hinder their effectiveness in consistently improving the protein expression of mRNA molecules. Firstly, the existing methods primarily rely on predefined sequence features, such as CAI, to guide codon selection. However, these metrics often fail to correlate with the experimentally measured protein expression levels^14,15^, indicating that they do not accurately capture the complex factors governing mRNA translation. Secondly, the existing methods do not adequately account for the activity of translational regulators that influence mRNA translation, such as translation factors and RNA-binding proteins^16,17^. This lack of context-aware optimization may reduce the effectiveness of the optimized mRNA sequences in specific cellular environments. Furthermore, the existing methods explore a limited space of codon sequences due to computational constraints and the reliance on predefined rules. This restricted search space may prevent the discovery of novel and highly optimized sequences that could potentially yield significant improvements in protein expression.

Deep learning has achieved remarkable success in tasks such as image recognition, natural language processing, and protein structure prediction, where it has outperformed conventional algorithms by learning complex patterns and relationships from vast amounts of data^18,19^. In the context of mRNA codon optimization, a deep learning approach may enable the model to capture the complex interplay between codon usage and cellular context, without relying on predefined rules. Moreover, deep learning models can explore a vast sequence space and discover novel patterns that may not be apparent to human experts or accessible through traditional optimization methods^20^. This ability has been exemplified in the field of protein engineering, where deep learning has been used to design novel protein sequences with improved stability, binding affinity, and catalytic activity^21–23^. Recent advances in codon optimization research have seen the emergence of deep learning-based algorithms, particularly large language models (LLMs) trained on cross-species nucleotide sequences. These models have been implemented for predictive modeling of mRNA translation efficiency and degradation kinetics^24,25^. Nevertheless, there persists an urgent demand for developing a rigorously validated optimization framework to improve mRNA-encoded protein expression specifically tailored for therapeutic applications.

Massive parallel reporter assays (MPRA) are commonly used to study the effects of regulatory sequences on gene expression^26^. However, it is not suitable for optimizing coding sequences due to the short sequence limitation, which is generally less than 300 base pairs, for high-throughput DNA synthesis. Additionally, MPRA experiments often rely on artificial reporter constructs and may not fully recapitulate the complex regulatory landscape of endogenous mRNA molecules. Ribosome profiling sequencing (Ribo-seq) is a powerful experimental technique that provides a snapshot of actively translating ribosomes on mRNA molecules^27,28^, where the translation level of an mRNA can be derived from the reads per kilobase per million (RPKM) of Ribo-seq. Recent studies have leveraged Ribo-seq to develop translation-focused deep learning models^29–31^. For instance, RiboNN predicts mRNA translation efficiency in mammalian cells by integrating mRNA sequences with ribosome profiling data, revealing translation-stability regulatory mechanisms^32^. However, the field critically requires a rational design framework that translates data-derived translational signatures into codon optimization strategies, enabling high-throughput exploration of sequence space and generation of optimized mRNA constructs for therapeutic development.

In this study, we present RiboDecode, a deep learning model for mRNA codon optimization that enhances mRNA translation by directly learning complex relationship of mRNA codon sequences to their translation level from large-scale Ribo-seq data. Our prediction model demonstrated robust performance, while analysis of RiboDecode’s optimization strategies revealed a complex interplay between sequence characteristics and translation. *In vitro* experiments showed significantly increased in mRNA translation and protein expression, outperforming past methods. RiboDecode also considered cellular context, and maintained robust performance across unmodified, m^1^Ψ-modified, and circular mRNA formats. *In vivo*, optimized influenza virus hemagglutinin (HA) mRNA induced approximately ten times stronger neutralizing antibody responses in mice, while optimized nerve growth factor (NGF) mRNA achieved equivalent neuroprotection of retinal ganglion cells at one-fifth the dose in an optic nerve crush mouse model. This data-driven approach to codon optimization advances our understanding of mRNA translation and facilitates the development of more effective mRNA therapeutics.

## RESULTS

### RiboDecode is a Deep Learning Framework for mRNA Codon Optimization

RiboDecode is a deep learning-based framework for optimizing mRNA codon sequences. It integrates three components: a translation prediction model, an MFE prediction model, and a codon optimizer that explores and optimizes codon choices guided by the prediction models (Figure 1a).

**Figure 1.**
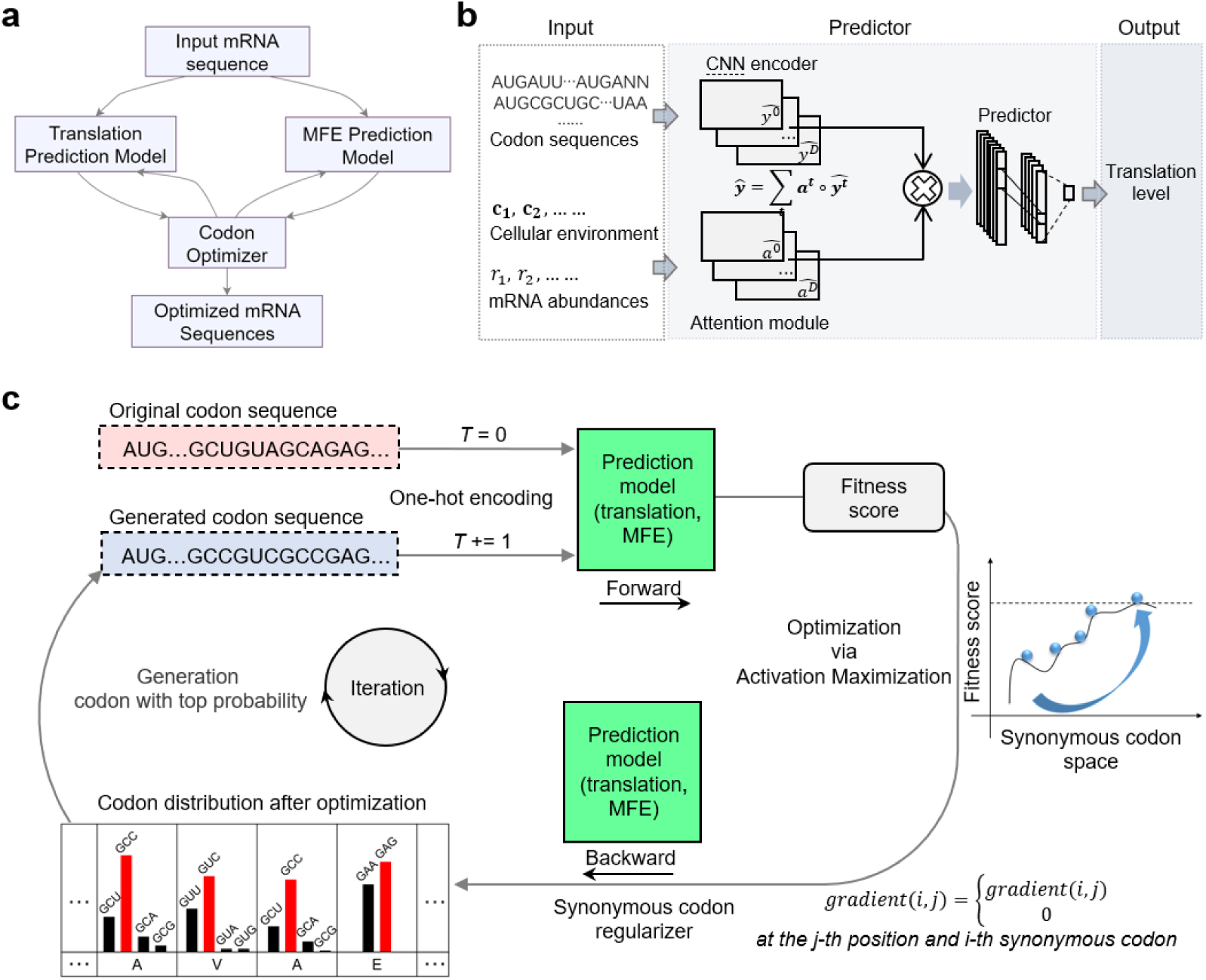
Predictive and Generative Optimization of RiboDecode. **a.** RiboDecode contains three main components, a codon optimizer, a translation prediction model and an MFE prediction model. **b.** The framework of the prediction model for translation. The input includes codon sequences in one-hot encoding, corresponding mRNA abundances and, the cellular environment which is presented by vectors of gene expression profiles from RNA-seq. The model integrates these inputs using CNNs. From the fused representations, a convolutional neural network extracts features and outputs the predicted translation level of mRNA (see Methods). **c.** Iterative optimization of codon sequences. RiboDecode predicts fitness of an original sequence (T=0), then uses activation maximization to generate optimized synonymous variants (T+=1). A synonymous regularizer maintains amino acid sequence. This process iterates until peak fitness is achieved.

The translation prediction model estimates the translation level of a given codon sequence by learning the translational expression of diverse mRNA sequences from Ribo-seq experiments (Figures 1b and S1, Methods). In contrast to previous tools that rely on optimizing predefined features such as CAI, our deep learning model automatically extracts relevant features by training on 320 paired Ribo-seq and RNA sequencing (RNA-seq) datasets from 24 different human tissues and cell lines, encompassing translation measurements of over 10,000 mRNAs per dataset (Supplementary file 1 and Methods)^33,34^. In addition, the model incorporates not only codon sequences but also mRNA abundances and cellular context that is presented by gene expression profiles from RNA-seq (Methods). This approach enables the prediction of mRNA translation by jointly considering these important factors influencing translation.

To address mRNA stability, we developed an MFE prediction model. Current MFE prediction tools, such as RNAfold^35^ and Linearfold^36^, use dynamic programming. However, these methods are non-differentiable and thus incompatible with our codon optimizer described below. Our MFE model employs a deep neural network architecture and undergoes an iterative optimization process, to simultaneously improve its predictive capability and optimize sequences for lower MFE values (Figure S2, Methods).

The codon optimizer of RiboDecode begins with the original codon sequence of a given protein (Figure 1c). The prediction models then predict a fitness score for this sequence. Using a gradient ascent optimization approach based on activation maximization (AM)^37^, the optimizer adjusts the codon distribution to maximize the fitness score (Figure S3). A synonymous codon regularizer ensures that only synonymous codons encoding the same amino acids as the original sequence are considered, preserving the protein’s amino acid sequence. Through iterative cycles of sequence generation, prediction, and optimization, the system produces codon sequences with improved properties. RiboDecode can optimize mRNA translation, stability or both, by interfacing with both the translation and MFE models. This uses a parameter, *w*: *w*=0 optimizes translation only, *w*=1 optimizes MFE only, and a value 0<*w*<1 jointly optimizes both (Methods).

By combining data-driven predictions with high-throughput sequence generation, RiboDecode overcomes limitations of conventional heuristic approaches. It enables the exploration of a vast mRNA codon space, potentially uncovering optimized sequences.

### Evaluation of Translation Prediction Model

We first evaluated the RiboDecode’s performance and generalizability using three cross-validation datasets: “unseen genes”, “unseen environments”, and “unseen genes and environments”, which represented unseen genes and unseen cell types during training (Figure S4, Methods). The model achieved a coefficient of determination (R^2^) of 0.81, 0.89, and 0.81 for the three datasets, respectively (Figures 2a), indicating its robustness and ability to generalize.

**Figure 2.**
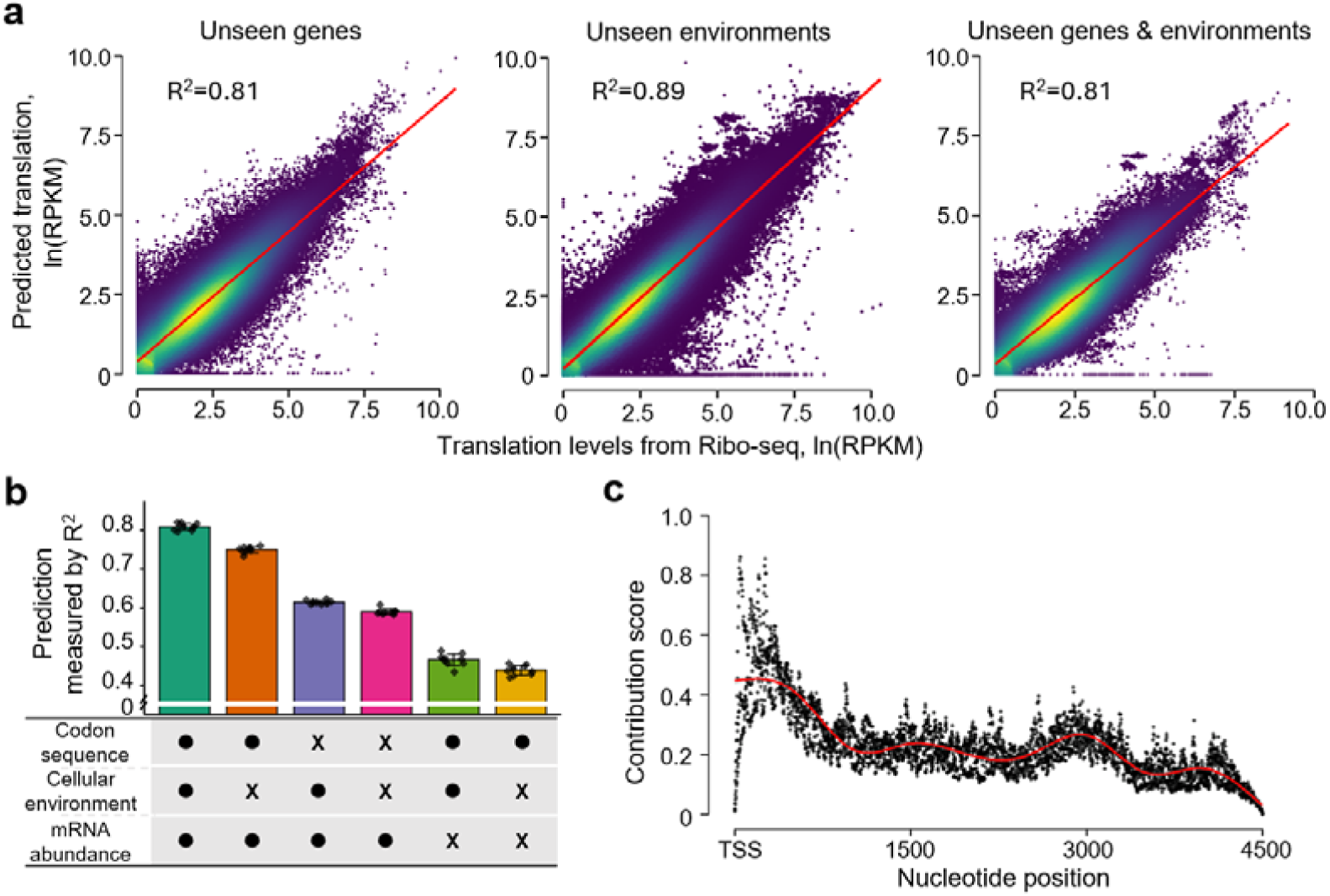
Evaluation of the Translation Prediction Model. **a.** Experimentally measured translation levels by Ribo-seq versus predicted translation levels in the three validation datasets. Red lines denote the linear fit. The translation levels from Ribo-seq were ln-transformed (see Methods). **b.** Ablation analysis shows the contributions of the three inputs to the prediction model. The table below shows the ablation status of the inputs, with dots and crosses representing the presence and absence of corresponding elements, respectively. The error bars denote standard deviation. **c.** The importance of each nucleotide position for the translation prediction. The x-axis represents the nucleotide position from the TSS (translation starting site). Integrated Gradients attribution method was used to obtain the importance score for each nucleotide position (black dots). The red line denotes the local polynomial regression fit.

To understand the relative importance of the three model inputs, we performed ablation analysis, revealing that mRNA abundances were the most important contributor to the prediction of translation (Figure 2b, Table S2), in agreement with an early study of yeast translation that found that the most predictive variable for translation was the mRNA expression of the gene^28^. The incorporation of codon sequences lifted the R^2^ by 0.15, and further inclusion of cellular environment improved the R^2^ by 0.06. The ablation analysis demonstrated that all the inputs contributed to predicting mRNA translation.

We next investigated whether our model captured complex sequence features beyond common translation-related metrics. While our model learned relevant sequence features directly from the raw codon sequences, we tried to include common translation-related sequence metrics including CAI, MFE, and codon frequencies as additional model inputs and found these metrics did not improve prediction accuracy (Table S1). This suggested that the model could capture the sequence patterns that were predictive of translation, beyond these sequence metrics.

We explored alternative approaches to incorporating cellular context information. We directly incorporated the meta information of Ribo-seq datasets into the model, including cell types and experimental conditions and found it did not improve the performance (Table S1). This indicated that the gene expression profiles used in the model were an effective proxy to capture the relevant cellular environment influencing mRNA translation.

Finally, we investigated the positional importance of coding sequences in translation prediction. We analyzed the importance of each nucleotide position for the model’s prediction (Methods). The results showed that the coding sequences close to the translation start site (TSS) were more important (Figure 2c). This is consistent with a general knowledge that codons near TSS have a greater impact on protein synthesis, by influencing translation initiation^8^.

Overall, the data-driven approach of RiboDecode enabled robust predictive capabilities with biological relevance by learning important sequence patterns directly from the Ribo-seq data.

### RiboDecode’s Optimization Strategies for Enhanced mRNA Translation

Having established the efficacy of our translation prediction model, we next explored how this model could be leveraged to generate sequences with enhanced translation potential. We first generated codon sequences of Gaussia luciferase (Gluc) (Figure S5). T-distributed stochastic neighbor embedding (t-SNE) indicated that the model established an association between the sequence space and translation levels (similar to Ribo-seq-derived RPKM values). The red area in the upper right showed that a wide space of high translation sequences was explored (Figure 3a). We next explored how RiboDecode optimized translation and stability independently or jointly. A widely used Gluc sequence was used as a reference for comparison, which had a predicted translation level of 5.9 and an MFE value of −216 (Figure 3b). By optimizing the sequence for translation (*w*=0), the predicted translation level increased to around 25. On the other hand, codon sequences optimized for MFE (*w*=1) reduced the MFE from −150 kcal/mol to around −350 kcal/mol, with a similar translation level to the reference. With joint optimization (0<*w*<1), RiboDecode explored a wider sequence space, achieving both enhanced translation and reduced MFE. Moreover, RiboDecode-generated sequences spanned a broader embedding space compared to those produced by Ribotree, CDSfold, and LinearDesign (Figure S6, see Methods), suggesting enhanced sequence diversity. Furthermore, designing Gluc codons for different cell lines showed that the generated codons had distinct sequence patterns for different cellular contexts, reflecting differences in cellular environment (Figure S7).

**Figure 3.**
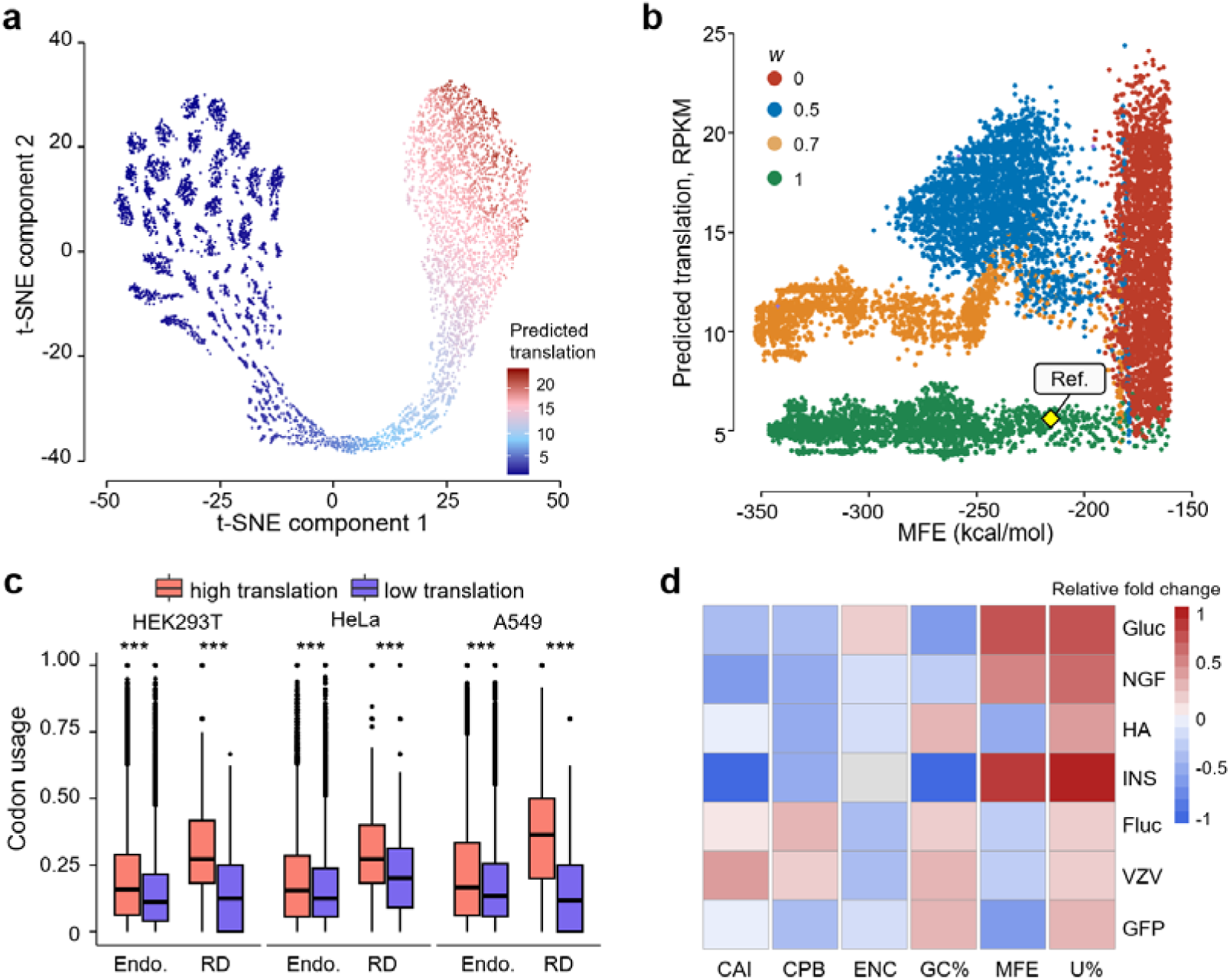
Strategies of Enhanced Translation in Generated Sequences. **a.** Generation of Gluc codon sequences with low translation level (the upper left area) and high translation level (the upper right area) (*w*=0, Figure S5). T-SNE of codon sequences is shown. Each dot represents one sequence, and the color represents the predicted translation level. **b.** Generation and optimization of Gluc codon sequences using different *w* of 0, 0.5, 0.7, and 1. Each dot represents one sequence, positioned in its predicted translation level (y-axis) and MFE (x-axis). The position of the reference sequence (MF882921.1) is shown. **c.** Codons that appeared more frequently in highly translated endogenous sequences were also used more often in highly translated Gluc sequences generated by RiboDecode. “RD”: RiboDecode-generated sequences, “Endo.”: endogenous genes. (***: *p*<0.001, t-test). **d.** Changes of sequence features of optimized sequences compared to the unoptimized, for different mRNAs. For each column (feature), cells in red represent that the values were higher in optimized sequences than unoptimized ones while cells in blue represent the opposite. The difference of ENC for INS shows no significance (the cell in gray). Abbreviations: GFP: green fluorescent protein, Fluc: firefly luciferase, INS: insulin, VZV: varicella zoster virus glycoprotein E, and HA: influenza A hemagglutinin.

To understand RiboDecode’s optimization strategy, we analyzed codon usage patterns between generated sequences with enhanced and reduced translation, as well as between high- and low-translated endogenous sequences. We found that codons preferentially used in highly translated endogenous sequences were also favored in RiboDecode-generated sequences with enhanced translation. Notably, the differences in codon usage between RiboDecode’s enhanced and reduced translation sequences were more pronounced than the differences observed in endogenous sequences (Figures 3c, Methods). To assess the generalizability of these findings, we extended our analysis to multiple genes across various cell types. Consistently, we observed the same pattern of biased codon usage in all the cases (Figure S8). This suggests that RiboDecode not only mimics but amplifies the codon usage patterns of efficiently translated endogenous mRNAs, potentially leading to even greater improvements in translation.

We next examined how RiboDecode utilized sequence features during generation and optimization. Analysis of sequence features across different mRNAs revealed complex and variable relationships with translation (Figures 3d and S9, Methods). Notably, highly translated mRNAs generally showed an increase in uridine content (U%), which may reduce secondary structure formation and facilitate smoother ribosome movement during translation^7^. Additionally, these mRNAs mostly exhibited a decrease in Effective Number of Codons (ENC), suggesting a selection against rare or inefficient codon pairs to enhance translation^38^. Variations in CAI, Codon Pair Bias (CPB), GC content (GC%) and MFE across different mRNAs suggested that while these features could influence translation, their impact might be more mRNA- or context-dependent, due to complex sequence or structure feature changes.

In short, these findings highlight RiboDecode’s ability to capture complex sequence-translation relationships, offering a sophisticated approach to mRNA optimization that goes beyond traditional codon optimization metrics.

### Experimental Validation Demonstrates RiboDecode’s Versatility and Efficacy

While our *in silico* analyses demonstrated the potential of RiboDecode to optimize codon sequences, we next sought to validate these findings experimentally. We first validated RiboDecode’s ability to optimize codon sequences for enhanced protein expression. For Gluc, protein expression levels of the RiboDecode-optimized sequences surpassed the reference and were more than twice as high as that of the best-performing sequences designed by LinearDesign (p-value=0.019, one-sided Mann-Whitney U test, Figure 4a and S17a, Table S3). The predicted translational levels showed a positive correlation with the experimentally measured protein levels (Correlation coefficient=0.71, p-value=0.077, Pearson’s correlation, Figure S10). However, the p value is marginal, potentially due to the small number of constructs tested. Further validation with larger datasets is needed. In contrast, the CAI showed negative correlation with the experimental measurements (Correlation coefficient=−0.15), indicating that CAI is not a reliable predictor of protein expression levels and ineffective optimization strategy in this context. We noticed that the two RiboDecode-designed mRNAs (RD1 and RD2) had the best performance which also had the highest MFE values of around −200. In contrast, the LinearDesign sequences had MFE values of −350 and −300. To rule out the increased protein production was associated with higher MFE value, we tested additional LinearDesign sequences with higher MFEs (−266.8 and −246.70 kcal/mol, respectively). Compared to four LinearDesign sequences, RiboDecode-designed sequences outperformed (Figure S11).

**Figure 4.**
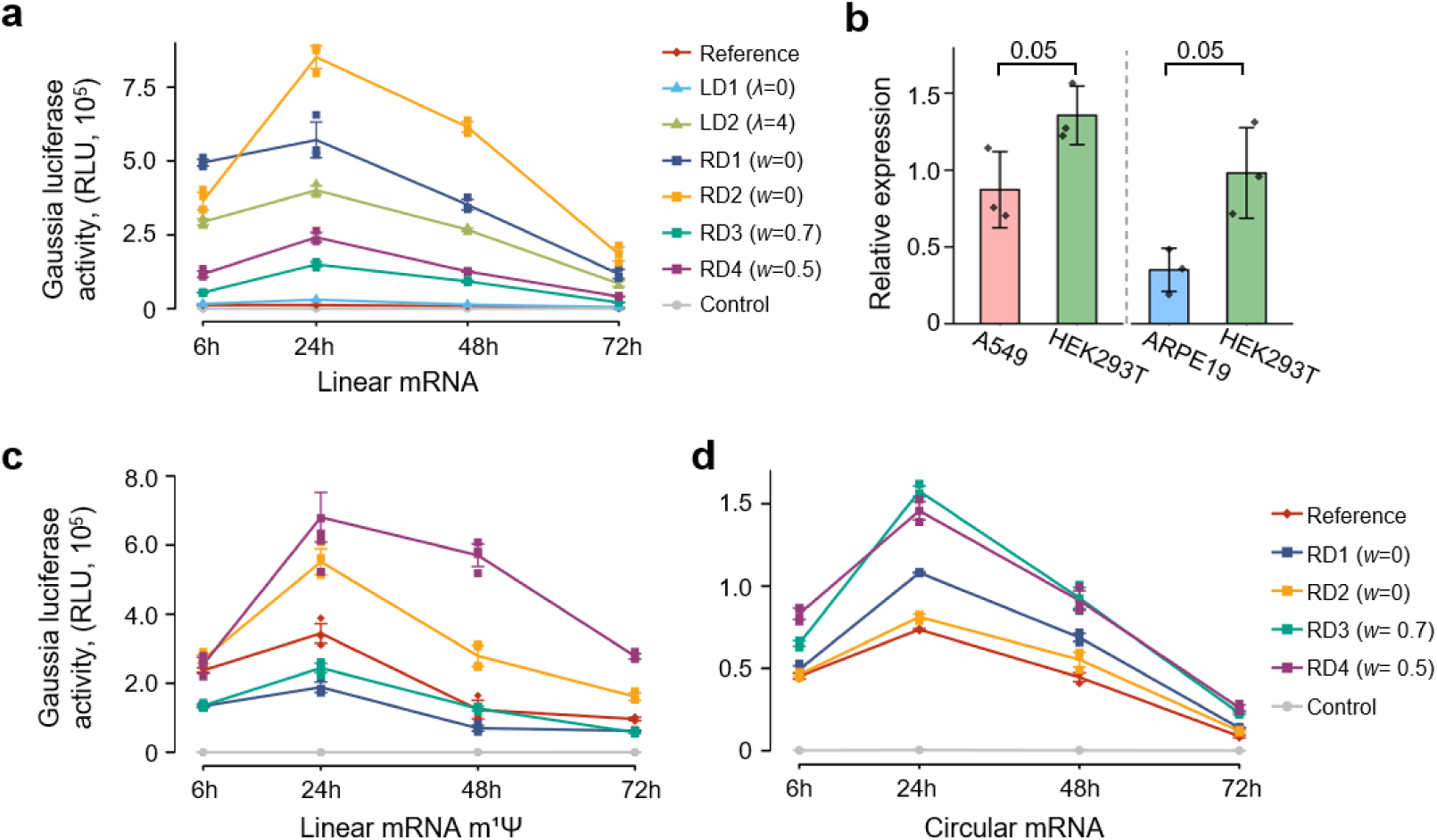
Robustness of Optimization across Unmodified, Modified, and Circular mRNAs. **a.** Protein expression of Gluc was measured by fluorescence intensity. RD sequences were designed by RiboDecode, with *w* parameter indicated in parentheses. LD sequences were designed by LinearDesign, with λ parameter indicated in parentheses. “RLU”: relative light units. The error bars denote standard deviation. **b.** Relative expression of experimentally measured protein expression values of mRNA variants designed by RiboDecode at 24 hours in different cells. The error bars denote standard deviation. One-sided Wilcoxon test was used to calculate p-values shown in the figure. **c, d.** Protein expression of generated Gluc codon variants in (c) the linear mRNA form with m^1^Ψ modification and (d) the circular mRNA form. The error bars denote standard deviation.

We additionally optimized another commonly used reporter gene, firefly luciferase (Fluc). Seven sequences, including four RiboDecode-optimized (RD1-RD4), two LinearDesign-optimized (LD1, LD2) and a WT, were transfected into HEK293T cells, and activity was measured over 72 hours (Figure S12, Table S6). All optimized sequences significantly outperformed the WT. The LD1 (CAI of 0.766) and LD2 (CAI=0.952) yielded the increased expression with about 7-and 41-fold changes, respectively. RiboDecode sequences (CAI of ∼0.71-0.73) also achieved substantial improvements with 6- to 16-fold over the WT. The superior performance of the LD2 sequence with a high CAI value here—contrasting with results for Gluc—underscores that the optimal codon strategy is gene-specific. This highlights the need for context-dependent approaches like RiboDecode that capture complex features beyond single metrics like CAI.

We next evaluated RiboDecode’s ability to design for cellular context by optimizing Gluc mRNA for preferential expression in HEK293T cells over A549 and ARPE19 cells. The designed variants successfully exhibited the intended higher expression in HEK293T compared to both cell lines (Figure 4b, Table S4). Predicted expression ratios favoring HEK293T closely matched experimental results in the comparison against A549 (∼1.7-fold predicted vs. ∼1.55-fold experimental). Preferential expression was also achieved against ARPE19 (Figure 4b, Table S5), although the experimental fold-change (∼2.8-fold) doubled the prediction (∼1.4-fold). These results confirm RiboDecode’s capability for context-aware design, while the discrepancy in the ARPE19 comparison indicates potential for refining the model to better capture quantitative differences across diverse cellular environments.

Modified mRNAs, such as those with 1-methylpseudouridine (m^1^Ψ) modifications, and circular RNAs are used in mRNA therapy instead of unmodified mRNAs due to their improved stability and reduced immunogenicity^3,7,39^. We therefore assessed the effectiveness of RiboDecode in enhancing translation in these alternative mRNA forms. Among the four codon variants, m^1^Ψ-modified RD2 and RD4 showed higher protein expression levels compared to the reference, with up to a 4.6-fold higher expression at 48 hours post-transfection (Figure 4c and S17b). Moreover, all four RiboDecode-generated codon variants in the circular form outperformed the reference (Figure 4d and S17c). These results demonstrate that RiboDecode optimization enhances protein production in both m^1^Ψ-modified and circular mRNAs, illustrating its reliability and versatility.

These experimental validations demonstrate RiboDecode’s ability to significantly enhance protein expression, optimize in specific cell types, and improve translation across various mRNA forms, highlighting its potential as a powerful tool for mRNA therapeutic development.

### RiboDecode Enhances Immunogenicity of mRNA-based Influenza Vaccines

Having established the robustness of our optimization approach, we next aimed to demonstrate its practical application in the development of mRNA-based vaccines. Influenza A viruses are responsible for causing respiratory infections, leading to annual epidemics that result in millions of human infections worldwide^40^. HA, a glycoprotein found on the surface of influenza A viruses, plays a crucial role in the viral infection process and is the primary target for the development of influenza vaccines. Although most of the vaccines were developed using inactivated influenza viruses, mRNA-based influenza vaccines are currently actively developed^41^.

To enhance the expression of HA and potentially improve the efficacy of HA-based vaccines, we optimized the HA coding sequence. Three out of four RiboDecode-optimized HA sequences and two LinearDesign-optimized sequences showed higher *in vitro* protein expression compared to the WT (Figures 5a, Table S7). Particularly, RD3 showed approximately 6-fold increase compared to the WT and LinearDesign-optimized sequences. In addition, RD3 exhibited considerably higher expression levels compared to the WT sequence in both m^1^Ψ-modified and circular mRNA forms (Figures 5b and 5c). These results again highlight the robustness and versatility of the RiboDecode-optimized sequence.

**Figure 5.**
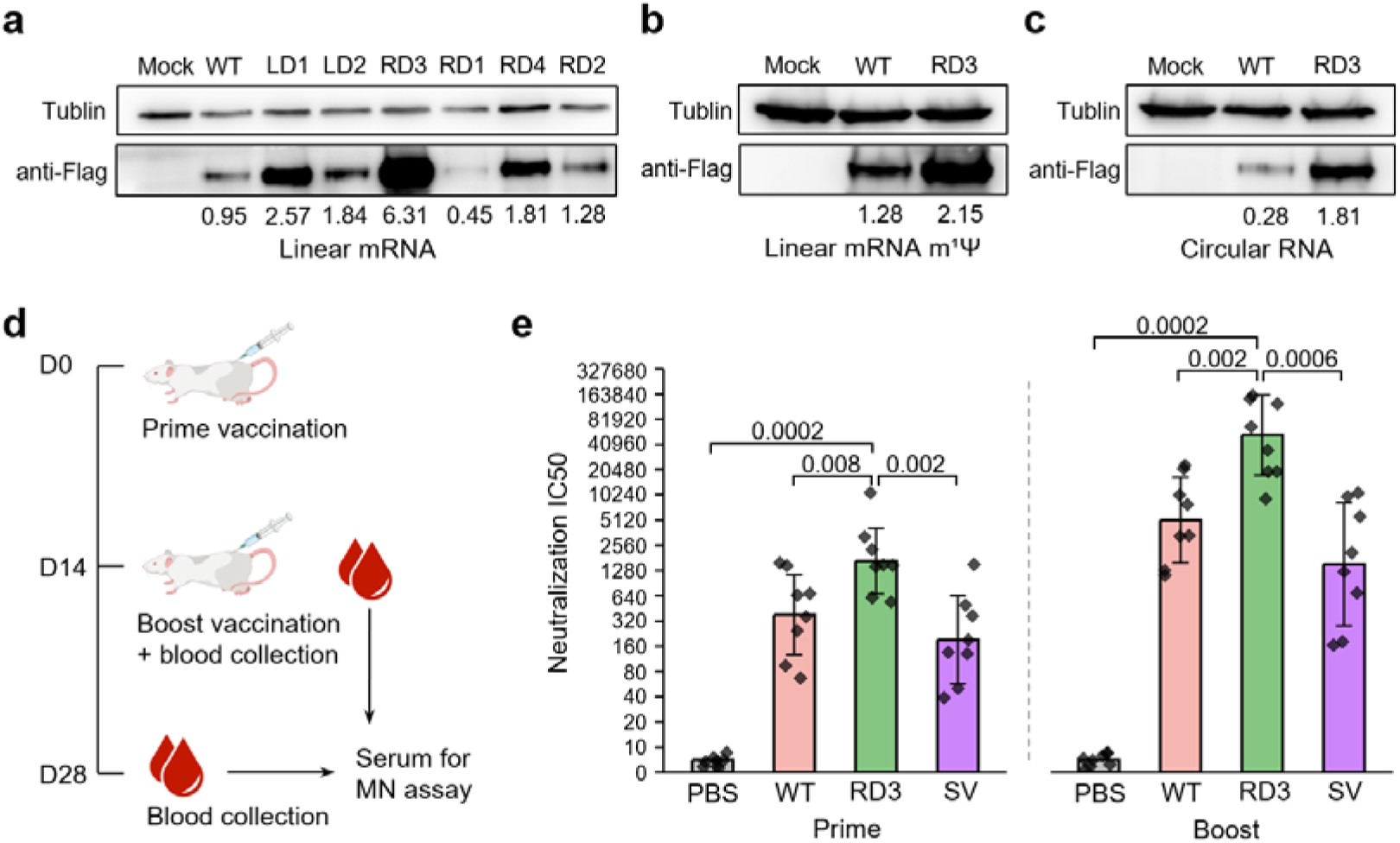
More Effective mRNA-based Influenza Vaccines Through Codon Optimization. **a.** Western blot analysis shows protein expression of the HA variants in HEK293T cells 24 hours after transfection. RD sequences were designed by RiboDecode (*w* parameter were set to 0, 0, 0.7, and 0.5 for RD1-4). LD sequences were designed by LinearDesign (λ parameter were set to 0 and 4 for LD1 and LD2). The expression values were quantified using GelAnalyzer. **b, c.** Protein expression RD3 in (b) the linear mRNA form with m^1^Ψ modification and (c) the circular mRNA form. **d.** HA mRNA immunization and analysis: BALB/c mice were intramuscularly inoculated with two doses (10μg mRNA for each dose) with an interval of two weeks. The mouse serum was collected at 14 days and 28 days for MN assay. **e.** Levels of neutralizing antibodies against influenza viruses after prime and boost vaccination. IC50, half-maximal inhibitory concentration. PBS and split virus influenza vaccine (SV) were used as the negative and positive controls, respectively. One-sided Wilcox test was used to calculate p values shown in the figure. The error bars denote standard deviation.

We further assessed the *in vivo* immunogenicity induced by the optimized sequence for both the prime and boost responses, where split virus influenza vaccine (SV) was served as the positive control (Figure 5d, Methods). The RD3 sequence induced significantly stronger neutralizing antibody responses, measured by the micro-neutralization (MN) titers, compared to the WT sequence and SV. For the prime response, RD3 elicited significantly higher MN titers compared to WT, with approximately 4.4-fold increase (Figure 5e, mean MN titers: RD3=2,560, WT=580; p-value=0.008, one-sided Wilcoxon test). The difference was more pronounced for the boost response, with RD3 inducing a 9.6-fold increase in MN titers compared to WT (Figure 5e, mean MN titers: RD3=83,200, WT=8,640; p-value=0.002, one-sided Wilcoxon test). These results demonstrated that the RiboDecode-optimized sequence significantly enhanced both the initial and boosted immune responses. This dramatic improvement in immunogenicity underscores RiboDecode’s potential to enable more effective vaccines with lower doses.

### Enhanced Protein Expression and Therapeutic Efficacy with optimized NGF mRNA

Having demonstrated the efficacy of RiboDecode in optimizing mRNA for vaccine development, we next explored its potential in protein replacement therapy. We focused on NGF as a promising candidate for treating glaucoma, which is a leading cause of irreversible blindness^42^ and causes death of retinal ganglion cells (RGCs). Our recent study demonstrated that mRNA-based NGF therapy provided robust neuroprotection for RGCs in an optic nerve crush (ONC) mouse model^43^.

To improve the neuroprotection efficacy, we optimized the codon sequences of human NGF mRNA. The protein expression levels of three RiboDecode-designed sequences were more than 2-fold higher compared to that of the WT whereas the LinearDesign sequences did not show improvement (Figure 6a and S18a, Table S8). We further assessed the best performing sequence (RD3) in both m^1^Ψ-modified mRNA and circular mRNA forms. Notably, with m^1^Ψ-modification, RD3 achieved 8.4- and 9.8-fold higher protein levels compared to the WT at 24h and 48h, respectively (Figure 6b and S18b). With mRNA circulation, RD3 also achieved a more than 2-fold higher expression than the WT at both 24h and 48h (Figure 6c and S18c). These results again demonstrated the robustness of RiboDecode-optimized sequences across different mRNA forms.

**Figure 6.**
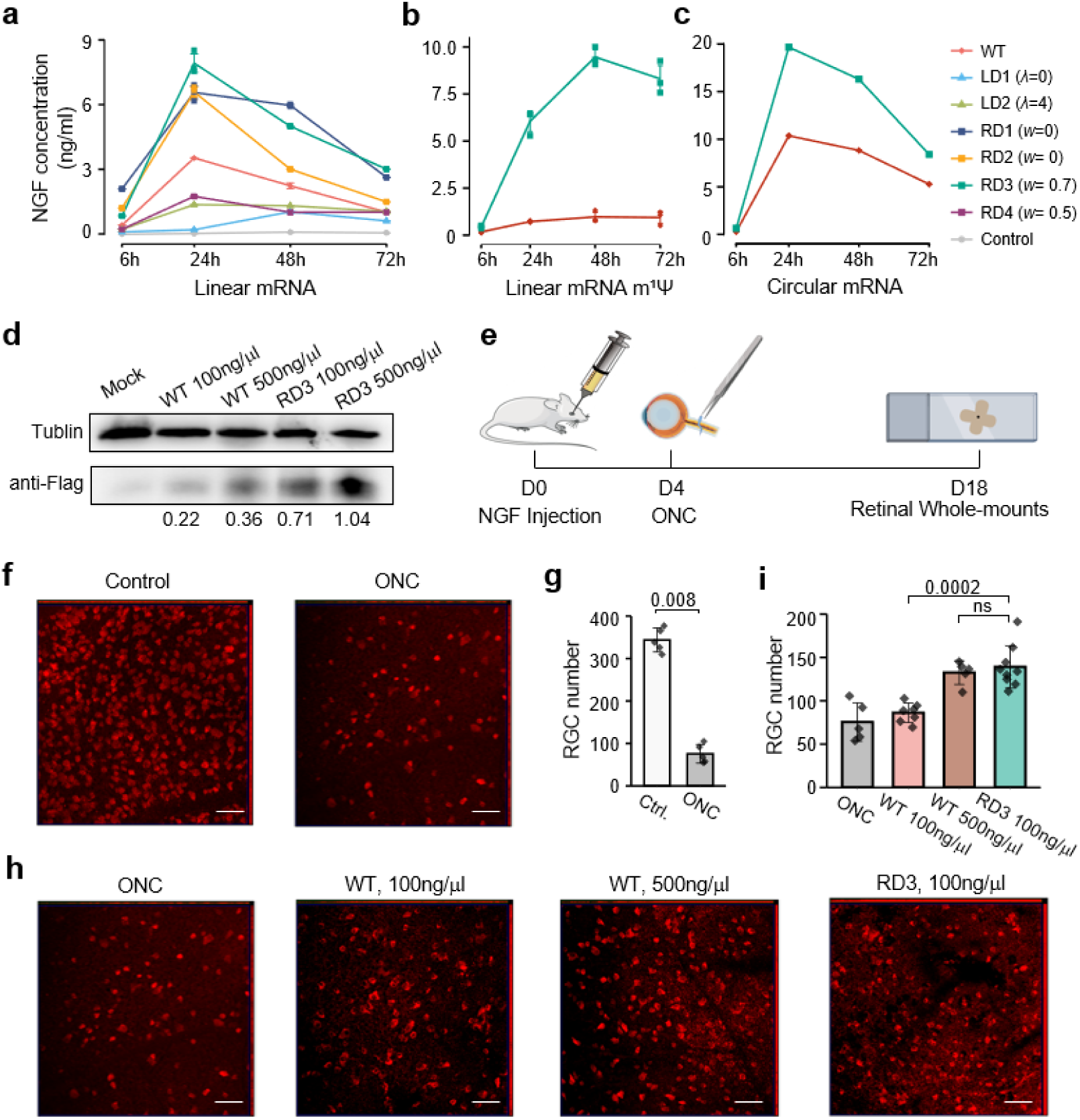
Enhanced Protein Expression and Therapeutic Efficacy with Optimized NGF mRNA. **a.** NGF protein expression in HEK293T cells, measured by ELISA. RD sequences were designed by RiboDecode, with *w* parameter indicated in parentheses. LD sequences were designed by LinearDesign. The error bars denote standard deviation. **b, c.** Protein expression levels of RD3 in m^1^ Ψ-modified (b) and circular (c) mRNA formats. **d.** *In vivo* protein expression of RD3. The m¹Ψ-modified mRNAs were injected into mouse retinas, and protein levels were measured by western blot 48 hours post-injection. The expression values were quantified using GelAnalyzer. **e.** Timeline of NGF mRNA therapy in ONC model: At Day 0 (D0), the m¹Ψ-modified NGF mRNAs were injected into mouse retinas. At Day 4 (D4), the optic nerve was subjected to a physical crush injury. At Day 18 (D18), RGC numbers were quantified using immunofluorescence staining. **f, g.** RGC numbers measured by immunofluorescence staining in the ONC mouse were significantly reduced compared to that of the control (one-sided Wilcox test). Scale bar, 50μm. **h.** RGC numbers in the ONC mouse retina after injection of NGF m^1^ −modified mRNA with 100 and 500 ng/μl dosages. Scale bar, 50μm. **i**. The RGC number in mice treated with 100 ng/μl RD3 was similar to those treated with 500 ng/μl WT and significantly higher than those treated with 100 ng/μl WT. One-sided Wilcoxon test was used to calculate p-values shown in the figure; ns: *p*>0.05.

Based on its superior performance in initial *in vitro* tests, we selected RD3 for further *in vivo* studies. To evaluate the *in vivo* expression of optimized NGF mRNA, we intravitreally administered both the RD3 and WT sequences. Each mRNA was m1Ψ-modified and encapsulated within LNP and administered at two doses: 100 ng/μl and 500 ng/μl. The RD3 sequence demonstrated significantly higher NGF protein expression compared to the WT sequence at both doses. Remarkably, RD3 at 100 ng/μl achieved even slightly higher expression than WT at 500 ng/μl (Figure 6d).

We then investigated the therapeutic potential of optimized NGF mRNA using an ONC mouse model, which mimics RGC injury and resulted in significant RGC loss (Figures 6e-g). Treatment with NGF mRNA showed clear neuroprotective effects, preserving more RGCs after injury. Notably, mice treated with 100 ng/μl RD3 showed significantly higher RGC counts than those treated with the same dose of WT mRNA (Figures 6h and 6i, p-value=0.0002, one-sided Wilcoxon test). Moreover, these counts were comparable to those in mice treated with 500 ng/μl WT mRNA.

To sum, the optimized sequence exhibited superior protein expression both *in vitro* and *in vivo*, while maintaining therapeutic efficacy at one-fifth the dose of the unoptimized sequence. These results demonstrated the effectiveness of RiboDecode in optimizing NGF mRNA for the treatment of RGC injury.

## DISCUSSION

In this study, we present RiboDecode, a novel deep learning-based framework for mRNA codon optimization. The generative optimization framework, guided by the deep learning prediction model, enables the efficient exploration of the immense space of possible codon sequences. This allows RiboDecode to discover novel, highly optimized sequences that may not be accessible to traditional optimization methods. RiboDecode-optimized sequences demonstrate superior performance in various mRNA formats, including unmodified, m^1^Ψ-modified, and circular mRNAs, highlighting its broad applicability in the rapidly evolving field of mRNA therapeutics. *In vitro* and *in vivo* experiments using the optimized sequences of therapeutically relevant proteins show substantial enhancements in protein expression compared to the unoptimized sequences. These improvements further translate into increased therapeutic efficacy, as demonstrated by significantly enhanced immune responses to an optimized influenza vaccine and markedly improved RGC protection in mice with optic nerve injury.

While mRNA abundance serves as both an input feature and intrinsically linked to the Ribo-seq RPKM target variable, potentially explaining its high predictive weight in ablation analysis, the model demonstrated significant improvements in prediction accuracy by integrating codon sequences and cellular context. Importantly, RiboDecode’s *in vitro* and *in vivo* validation, where optimized sequences substantially enhanced protein expression and therapeutic efficacy, confirms its ability to optimize biologically meaningful translational signals rather than merely reflecting transcript abundance. The superior performance of RiboDecode may be attributed to several factors. Firstly, RiboDecode’s deep learning model learns directly from diverse nature sequences with translation measurements, enabling it to capture complex patterns of codon sequences for mRNAs with high translation level. Second, the model considered the cellular contexts of mRNA translation. Third, RiboDecode’s generative optimization framework allows it to explore a large sequence space and to discover novel, highly optimized sequences that may not be accessible to the traditional approaches.

These findings also align with the broader framework of gene expression regulation: while mRNA abundance is a prerequisite for protein synthesis (consistent with its predictive dominance), our approach directly targets translational control, a critical layer highlighted by Schwanhäusser *et al.*^44^, by leveraging Ribo-seq-derived codon usage and cellular context to maximize translational efficiency. Although the model excludes post-translational events (as Ribo-seq captures ribosome activity prior to protein maturation), the achieved gains in protein output and *in vivo* efficacy robustly validate translational optimization as a key determinant of functional protein levels, complementing established regulatory hierarchies.

The findings of our study have important implications for the field of mRNA therapeutics. Firstly, RiboDecode can generate and evaluate a vast number of novel codon combinations. This capability allows RiboDecode to optimize mRNA sequences beyond the limitations of evolutionary constraints, potentially uncovering more efficient codon usage patterns not found in natural transcripts. Second, by substantially increasing protein production, the optimized sequences can improve the potency and reduce the required dose of mRNA-based treatments, potentially mitigating side effects and enhancing patient outcomes. This is particularly relevant for applications such as protein replacement therapies, where achieving high levels of protein expression is crucial for therapeutic success. Third, RiboDecode’s robustness and versatility across different mRNA formats, including modified and circular mRNAs, expand the range of therapeutic applications for which it can be employed. As the field of mRNA therapeutics continues to evolve and new mRNA formats are developed to enhance stability, reduce immunogenicity, and improve delivery^39,45^, RiboDecode’s ability to optimize sequences for these diverse formats will be invaluable.

While our study demonstrates the significant potential of RiboDecode in optimizing mRNA codon sequences for enhanced mRNA translation and therapeutic efficacy, there are several future directions to explore. Firstly, we focused exclusively on optimizing the codon sequences while not explicitly modeling the 5’UTR. Recognizing the critical role of the 5’UTR in regulating translation initiation, our future work aims to expand RiboDecode to jointly optimize both UTRs and the codon sequence. Second, our results indicate that while our primary goal was to enhance translation through codon optimization, incorporating MFE optimization synergistically modulating mRNA secondary structure stability and translation efficiency. However, we observed the best *w* value appears context-dependent, suggesting a one-size-fits-all approach may be suboptimal. Consequently, we recommend that experimental designs include the testing of multiple *w* values to identify the optimal balance for each mRNA target. Further research into the determinants of the optimal *w* value will be critical to refine this strategy and could lead to more systematic approaches for integrating MFE into mRNA design. Third, our MFE model cannot predict MFEs for unseen mRNAs. For an unseen mRNA, the model must first train the sequences with MFEs labelled by RNAfold. A general MFE model should be developed and used in future. Finally, the model was trained exclusively on Ribo-seq data from endogenous, unmodified mRNAs. Although our results showed significant expression enhancements for optimized sequences in both m1Ψ-modified and circular forms compared to their respective controls, the relative fold-improvement compared to the unmodified format varied between constructs. Future iterations could potentially incorporate data from modified and circular transcripts to further refine optimization rules.

In conclusion, RiboDecode represents a paradigm shift from rule-based to data-driven mRNA optimization, potentially uncovering entirely new principles of efficient translation that were previously inaccessible. RiboDecode will provide a versatile tool for researchers to maximize the potential of mRNA-based therapeutics, paving the way for more effective treatments in various medical applications.

## MATERIALS AND METHODS

### Data collection and processing

#### Data preprocessing and filtering

We downloaded translation counts of Ribo-seq datasets from the RPFdb database^33,34,46^. The following steps were implemented for processing the Ribo-seq data in accordance with RPFdb: First, to prevent adapter interference in downstream analyses, the 3’ adapter sequences were manually extracted for each dataset from the original publications or the corresponding MultiQC^47^ outputs. Adapter sequences, if present at the ends of sequencing reads, were subsequently removed using Cutadapt (version 1.16)^48^. Next, to minimize rRNA and tRNA contamination, sequences corresponding to rRNA and tRNA were retrieved for each species from ENSEMBL^49^ and UCSC^50^ databases and removed post-mapping using Bowtie2^51^. Finally, to ensure the retention of high-quality ribosome-protected footprints, which exhibit a characteristic read-length distribution reflecting the size of a translating ribosome on the RNA, only footprints within the 25–34 nucleotide length range were retained after contaminant removal and alignment. The count tables were transformed to reads per kilobase per million (RPKM). Because some of the paired RNA-seq were not available in RPFdb, we reprocessed the RNA-seq datasets. The raw FASTQ files of RNA-seq were trimmed by sickle^52^ for adapter removal and quality control. Then, to filter out reads from tRNA or rRNA, we mapped the reads to human tRNA and rRNA reference sequences (hg38) using bowtie2^53^ (-L 20). The unmapped reads were then mapped to the human genome (GRCh38, gencode.v28, https://www.gencodegenes.org/) using STAR^54^ (2.7.4a). Finally, read counts for each gene were summarized by featureCounts^55^ (v2.0.1, −t exon). The expression counts of Ribo-seq and RNA-seq were ln(RPKM x 5+1) transformed. Genes with low expression (median RPKM<1) were filtered out. Finally, 11,725 coding genes from 320 samples with 24 cell types were included in this study (Supplementary file 1).

#### Cross-validation dataset preparation

Out of 11,725 genes in the Ribo-seq data, we randomly selected 1,173 genes (1/10 of the total genes) that were not included in the training datasets, as the “unseen genes” dataset. Out of 320 Ribo-seq datasets, we randomly selected 120 datasets whose cell types were not included in the training datasets, as the “unseen environments” dataset (Supplementary file 1). The “unseen genes and environments” dataset was also defined (Figure S4).

#### mRNA isoform selection and sequence encoding

To address the complexity of alternative splicing while managing computational feasibility, we utilized the major isoform of each gene as a representative for mRNA codon variants. This approach was necessitated by the inherent limitations of NGS data in accurately quantifying the proportions of individual isoforms. We defined the major isoform as the transcript with the highest expression level, estimated using RSEM^56^. In total, our dataset contained 60,255 different mRNA sequences.

### Translation Model Architecture

Translation is influenced by multiple factors, including codon sequences as the pivotal signals modulating translation^57^, trans-acting elements modulating the cellular environment of translation^58,59^, and mRNA abundance which provides more templates for translation^60,61^. To capture these interacting variables, a translation model was developed using a deep neural network architecture comprising 2 convolutional layers and 5 fully connected (FC) layers^62,63^. The model inputs are a codon sequence in one-hot encoding, a transcript abundance, and a vector of gene expression profiles from RNA-seq presenting cellular environment (Figure S1). The codon sequences were fixed to 4,500 base pairs (bp) starting from the translation start site, which covers over 98% of coding sequences, and those shorter than 4,500 bp were zero-padded on 3’ end. To process these inputs, 3 sequential FC layers extract features from the cellular environment vector, which are then concatenated with the transcript abundance to form an attention vector. In parallel, convolutional neural networks (CNNs) processes the one-hot encoded codon sequence, generating embeddings with four distinct feature sets. Subsequently, the averaged feature is encoded through the other convolutional layers and flattened. Finally, 2 FC layers yield the predicted translation level output. To enhance model robustness and prevent overfitting, batch normalization^64^ and dropout^65^ techniques are employed after each layer, and max-pooling^66^ is utilized following the convolutional layers. For optimization, the AdamW optimizer^67^ is adopted in conjunction with the SmoothL1Loss loss function and ReLU activation functions^68^. Additionally, gradient clipping and learning rate decay strategies are implemented to mitigate gradient explosions and instability during training. The model was parameterized by a total of 15 hyper-parameters, as detailed in Table S9. The model was pretrained for 20 epochs before being used by the codon optimizer of RiboDecode. Model training and validation were assessed using the R-squared metric (R^2^). The best-performing model was selected based on the highest R^2^ value achieved on the validation set. The R^2^ metric, defined as R^2^ = 1 − (SS_Residual_/SS_Total_), quantifies the goodness of fit by comparing the sum of squared residuals (SS_Residual_) to the total sum of squares (SS_Total_). The PyTorch 1.12.0 framework was leveraged for the implementation of the deep model.

The translation model’s architecture, which uses a fixed 4,500 bp input, results in a high nominal parameter count. This is an artifact of flattening the zero-padded input sequences required for the majority of genes, which have a median length of only 1,200 nt. To mitigate the resulting risk of overfitting, we implemented a strong dropout regularization strategy (rate of 0.9) before the final layers. This high dropout rate forces the model to learn robust features rather than fitting to noise in the padded regions. A comparative study (Figure S15) confirms this strategy is essential: models with no or low dropout overfit immediately, whereas the 0.9 dropout rate eliminates overfitting and ensures stable, improving performance on all validation sets.

### MFE Model Architecture

The MFE prediction model was formed by a deep neural network architecture comprising 2 convolutional layers, 9 residual blocks derived from ResNet^69^, and 3 FC layers (Figure S2a). The model input is the one-hot encoded codon sequence, which is the same as the codon sequence used in the translation model. Each residual block consists of 2 convolutional layers with a residual connection. Initially, shallow features are extracted through 2 convolutional layers, followed by 5 residual blocks with max-pooling for deep feature extraction. Subsequently, 4 residual blocks without max-pooling are used to maintain spatial resolution. After the final residual block, features are flattened and fed into 3 FC layers to predict the MFE value. Batch normalization^64^ is applied after the first 2 convolutional layers, and dropout^65^ is introduced after the subsequent 2 residual blocks. The Fast Gradient Method (FGM)^70^ is integrated during training to enhance generalization by applying perturbations to input sequences. We utilized the AdamW optimizer^67^, SmoothL1Loss function, and LeakyReLU activations^71^. The final loss is defined as Loss = Loss_mfe_ + Loss_fgm_, where Loss_mfe_ is the SmoothL1Loss between predictions and RNAfold^35^ MFE values, and Loss_fgm_ is the SmoothL1Loss with added perturbations. The model is trained alongside optimization by the codon optimizer of RiboDecode using generated sequences as the training data, with performance evaluated using the R^2^ metric. Details of 15 hyperparameters are provided in Table S10. The PyTorch 1.12.0 framework and RNAfold 2.4.18 were used through Python interfaces. We evaluated the MFE values of mRNAs between our model and RNAfold. We found that our MFE value highly agreed with the one from RNAfold (Figure S13a), showing the reliability of our MFE model. To determine the optimal training set size, we compared our model trained on a compact set of 210,000 sequences against a model trained on an extended dataset of 11.5 million sequences (achieving a data-to-parameter ratio > 1). Both models achieved comparable optimization performance and yielded MFE predictions highly correlated with values from RNAfold (Figure S13). Given the similar performance, we selected the 210K training set for our framework, as it reduces computational time by approximately 75% without sacrificing model reliability or accuracy.

We evaluated our MFE optimization framework against existing MFE optimization methods, including the general-purpose, differentiable model (JAX-RNAfold) ^72^. Our analysis revealed that the high computational complexity of JAX-RNAfold severely limits its use to short sequences (under 600 nt on our V100 GPU with 32 GB video RAM), making it unsuitable for our primary goal of optimizing full-length human mRNAs. To perform a direct, head-to-head comparison, we used a compatible sequence (Gaussia luciferase, 558 nt). Our model achieved a lower (more favorable) MFE, completing the optimization faster and with substantially less GPU memory than JAX-RNAfold (Table S12). While LinearDesign is another powerful tool, it is not a differentiable model and is thus incompatible with our gradient-based optimization framework. Given the critical need for scalability to therapeutically relevant sequence lengths and the efficiency observed in direct comparison, we proceeded with our sequence-specific MFE optimization approach.

### Codon Optimizer Architecture

#### Fitness score

The fitness score combines mRNA translation level and MFE predicted by above models.

For translation level, the prediction of a mRNA sequence is conducted using the pre-trained translation model with the cellular environment and transcript abundance holding constant during the optimization. The loss function is designed as follows:

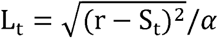

Here, S_t_ is the predicted translation level, r represents the desired output value of the translation model, and α is a constant based on S_t_.

For MFE, the loss function is formulated based on model parameters of the current epoch since the MFE model is trained alongside the optimization:

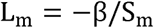

where, S_m_ is the predicted value of the current epoch and β is a constant set based on it.

The introduction of α and β is intended to balance the loss in the translation optimization and MFE optimization processes. The output range of the translation model (SC) typically lies between 0 and ∼100 (e.g., 0 to 25 for Gluc), while RNAfold-derived MFE values generally range between −100 and −1000 (e.g., −350 to −150 for Gluc). To ensure comparable magnitudes and numerical stability during optimization, we introduced scaling factors α and β to normalize both loss components. For most codon sequences exhibiting translation prediction values < 100 and MFE values > −1000 kcal/mol, α=β=100 provide appropriate normalization (Table S11). However, we recommend parameter adjustments under extreme value conditions: α should be increased to 1000 when translation predictions exceed 100, while β should be elevated to 1000 when MFE values fall below −1000 kcal/mol.

The final optimized loss function is defined as follows:

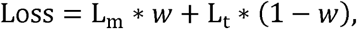

Where *w* is a constant ranging from 0 to 1. When *w* is set to 0, only the translation of mRNA is optimized. When *w* set to 1, only the MFE of mRNA is optimized. When *w* set to a constant value between 0 and 1, both models are optimized simultaneously, with the magnitude indicating the relative emphasis on optimizing each model.

#### Optimization process of the codon optimizer

The codon optimizer uses a gradient ascent optimization approach based on activation maximization (AM)^30^ to generate synonymous codon sequences with optimized fitness score. The optimization process involves the following steps:

1. Initial representation: the codon distribution is initially represented in a one-hot manner, where each position is assigned to the specific codon of the original codon sequence.
2. Optimization: the optimized codon distribution by AM becomes probabilistic, assigning a likelihood to each possible synonymous codon at every position, with the goal of maximizing the fitness score.
3. Regularization: during the optimization, a synonymous codon regularizer is used to ensure that the optimization process only adjusts the selection probabilities within the synonymous codons capable of encoding the same amino acids as in the original sequence. The regularizer applies a constraint on the codon distribution, T, given by a synonymous substitution mask matrix *W*:

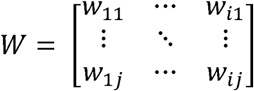

where w_ij_ signifies the selection of the j^th^ coding category at the i^th^ position, with value 1 for a synonymous codon, and 0 otherwise. Here, i = 1, 2, …, L and j = 1, 2, …, C. Subsequently, regularization is performed on T:

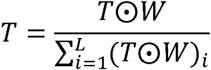 The maximum index values are then converted to a one-hot representation to obtain T_one-hot_:

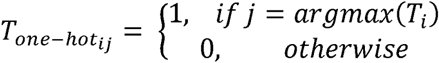
4. Sequence generation: after the gradient updates, a new codon sequence is generated by selecting codons with the highest probabilities.
5. Iteration: This sequence then re-enters the codon optimizer for further rounds of optimization.

We employed the Adam optimizer and utilized a learning rate decay strategy at different training stages. The optimization process was regularized with a weight decay of 1×10^−4^. The learning rate was initialized at 5×10^−4^ and decayed by a factor of 10 when the generated data reached 1/2, 3/4, 7/8, and 16/17 of the total 40,000 sequences for one epoch. The optimization process was conducted over 20 epochs. However, we observed that all genes converged to maximum translation levels before the 7th epoch. Consequently, a total of 280,000 sequences were utilized to plot the progression of the generation.

#### Integrative training of MFE model

To train the MFE model, we adopted an active learning strategy that integrates model training with sequence optimization by the codon optimizer. This approach uses the RNAfold tool^31^ for labeling and involves simultaneous training, optimization, and sequence generation. The process consists of four interconnected steps that run concurrently with the sequence optimization:

Initial sampling: we begin by generating 20,000 sequences through random synonymous substitutions (up to 10% of codons) of the original sequence.

Initial training: these 20,000 sequences are input into both the MFE model for predictions and RNAfold for ground truth MFE labeling.

Sequence generation: utilizing the trained model of current epoch, new sequences are generated by the codon optimizer of RiboDecode. This pool is sorted by predicted MFE, with lower values being better. The top 480 sequences are selected as generated sequences. The top sequence undergoes two operations: random replacement (up to 10% of codons) and distributed replacement according to its generating codon distribution. A total of 10,000 sequences are generated through this process.

Model retraining: The generated sequences, along with those from distributed and random replacements, are used as input for MFE prediction and RNAfold labeling. The model is then retrained on this new data.

#### Model running time

The computational infrastructure employed for our model training and sequence optimization comprised a single NVIDIA Tesla V100 SXM2 GPU with 32 GB of memory and an Intel Xeon Gold 5218 CPU operating at 2.30 GHz.

The translation model required approximately 24 hours for training completion. For the Minimum Free Energy (MFE) model, training and optimization were conducted concurrently. The duration of this process varied based on input sequence length, ranging from 1 hour to 3 hours. Upon completion of model training, the optimization phase was executed, which took about 1 hour for any sequences.

#### Optimized codon sequences screening strategy for experimental validation

To select optimal codon sequences for experimental validation, we employed a multi-step screening process for each gene:

1. Sequence Generation: We generated three rounds of candidate sequences using different optimization strategies: a) Translation-only optimization (*w*=0) b) Joint optimization with moderate MFE consideration (*w*=0.5) c) Joint optimization with stronger MFE consideration (*w*=0.7).
2. Initial Filtering: For each round, we filtered out potentially over-optimized sequences by retaining only those variants with a predicted translation level below the 90th percentile of all generated variants. This step helps avoid unreliable over-optimization that might not translate to real-world performance.
3. Selection Criteria: From the filtered sequences in each round, we selected candidate sequences based on two criteria: a) High predicted translation level b) Low MFE value.
4. Final Selection: We recommend selecting one or more candidates from each of the three optimization rounds (*w*=0, *w*=0.5, *w*=0.7) for experimental validation. This ensures a diverse set of optimized sequences, balancing pure translation optimization with different levels of MFE consideration.

### Model evaluation and analysis

#### Translation model evaluation

To evaluate the importance of each input component, we independently trained multiple translation models on the cellular environment vector, transcript abundance, and codon sequence and their combinations^20^. During training, the hyperparameter settings and dataset partitioning were consistent for each model. We replaced the input to be ablated with a zero tensor of the same shape and independently observed the impact of each input component on the final translation prediction.

When evaluating the performance of the model with CAI, MFE and cellular information as additional inputs, these new features were concatenated into the attention vector of cellular environment features and transcript abundance for mRNA translation prediction.

Furthermore, potential associations between genes in the unseen dataset and those in the seen dataset may lead to overestimation of model performance. To evaluate this potential issue, we obtained gene family annotations from the HUGO Gene Nomenclature Committee (HGNC; https://www.genenames.org/) and excluded genes in the unseen dataset that belonged to the same gene families as those in the seen dataset (resulting in the removal of 130 genes). The results demonstrated only a marginal decrease in prediction accuracy on the “unseen gene test set” (with R² decreasing from 0.81329 to 0.81295). Therefore, although some of the genes in the unseen dataset are related to those in the seen dataset, the performance of the model was not overestimated.

#### Nucleotide position contribution for translation prediction

To evaluate the importance of each nucleotide at different positions in the codon sequence, we explored the attribution of translation model predictions to their input features. Here, we implemented the attribution method of Integrated Gradients^73^, obtaining an importance score for each nucleotide position. This method combines the implementation invariance of gradients with the sensitivity of techniques such as LRP^74^ or DeepLift^75^. Firstly, we determined a vector with all features set to zero as the baseline value. Then, we linearly interpolated the input features from the baseline value to the actual input values, with these intermediate values representing different strengths or combinations of features. Subsequently, for each interpolated input, we calculated the gradient of the model output relative to that input, and multiplied the gradient at each interpolation point with the difference between the input feature value and the baseline value, obtaining the contribution of that feature at each interpolation point. Finally, we weighted and summed the contribution values at all interpolation points to obtain the Integrated Gradient for that feature. For the final analytical outcomes and visualization, the importance score assigned to each nucleotide position was calculated as the mean of absolute values derived from non-padding regions.

#### Cellular environment and mRNA abundance for translation model

The default mRNA abundance level was set to 4.5 (ln-transformed RPKM×5+1, median transcript abundance; Figure S14). To evaluate robustness, we additionally tested input values of 3.82 and 5.19, corresponding to half and double the original RPKM expression level, respectively. Notably, Gluc codon sequence predictions remained highly consistent across these input variations, demonstrating the stability of our model.

When optimizing mRNA sequences using RiboDecode, we recommend using the RNA expression profile of the target or the most similar tissue or cell type through either experimental sequencing or publicly available datasets. To facilitate implementation, we have incorporated a user-defined environment input parameter into the software package, accompanied by comprehensive documentation and step-by-step instructions.

For example, to predict translation in HEK293T, the input of cellular environment was constructed as follows. First, the mRNA expression levels of genes representing environmental factors were obtained from RNA-seq data of untreated HEK293T cells. Then, the batch effect arising from different data sources was corrected with ComBat^76^. Finally, the mean value of the mRNA expression was taken as the cellular environment input.

#### Codon usage analysis

Codon sequence variants with enhanced and reduced predicted translation levels were generated in different cellular contexts, including HEK293T, HeLa, and A549. Codon sequences of endogenous genes with high (top 10%) and low (bottom 10%) translation level were selected from Ribo-seq data. We further evaluated the impact of varying expression thresholds (top/bottom 5%, 10%, and 20%, Figures S8 and S16), with consistent results observed across all cutoff values, except Gluc in A549 cells at the 5% threshold. Then, the codon usage of generated and endogenous sequences was calculated by the proportion of each codon among synonym codons. To get the codons that appeared more in high-translated sequences, we performed t-test (p-value < 0.05, after FDR adjustment for multiple-testing) on high and low-translated sequences for both endogenous and generated sets. Codons with the significant and greatest differences on codon usage in endogenous sequences were chosen as the top-10 codons.

#### Sequence feature analysis

To analyze the sequence features of RiboDecode-optimized sequences compared to non-optimized sequences, we followed these steps:

1. Sequence Generation: a) We randomly selected 2,000 RiboDecode-optimized codon sequences. b) We generated 2,000 non-optimized sequences by performing random synonymous codon substitutions on the unoptimized input sequence.
2. Feature Calculation: We calculated several sequence features for both sets of sequences, including Codon Adaptation Index (CAI), Codon Pair Bias (CPB), Effective Number of Codons (ENC), GC content (GC%), Minimum Free Energy (MFE) and Uracil content (U%).
3. Fold Change Calculation: For each feature, we calculated the fold change by dividing the median value of the optimized sequences by the median value of the non-optimized sequences.
4. Data Transformation: We ln-transformed the fold change values for each feature to normalize the distribution.
5. Data Scaling: The ln-transformed fold change values were scaled to a range of −1 to 1 for visualization purposes.
6. Visualization: The scaled values were used to create a heatmap representation of the feature changes (Figure 3d). For detailed distributions of each feature, refer to Figure S9.

#### Parameters used for sequence optimization

For Gluc, HA, and NGF sequence optimization, four sequences were designed by RiboDecode (RD1, RD2, RD3, and RD4: *w*=0, 0, 0.7 and 0.5, respectively), and two sequences were designed by LinearDesign (LD1 and LD2: λ=0 and 4). The WT sequence was also used as a reference.

To preliminarily evaluate mRNA optimization *in vitro*, we first performed codon optimization for HA and NGF for the HEK293T cellular environment and measured their protein expression in HEK293T cells (Figures 5a-c and 6a-c).

#### Comparative analysis of sequence space across design methods

We generated 1,000 full-length Gluc codon sequences using Ribotree, LinearDesign, CDS-fold, and RiboDecode, respectively. High-dimensional sequence embeddings were extracted using CodonBERT^24^, followed by t-SNE dimensionality reduction for visualization.

### mRNA preparation

#### Plasmid construction

The 5’homology sequence, IRES sequence, 3’homology sequence, E1/E2 sequence and protein coding sequence, were chemically synthesized, and cloned into the vector pUC57, which contains a T7 RNA polymerase promoter.

For linear mRNA, the plasmids contain the 5’UTR, protein coding region, 3’UTR and 105nt poly-A elements. A 3xflag tag was added after the coding region, in order to detect the protein expression by western blot. The UTR and coding sequences were listed in Supplementary file 2.

#### Linear mRNA production and modification

The linear mRNAs were produced using the HiScribe T7 High Yield RNA Synthesis Kit and capped with m7G(5’)ppp(5’)G RNA Cap Structure Analog (NEB, #S1404). Then, the RNA was column-purified.

For mRNA m^1^Ψ-modification, N1-Me-Pseudo UTP (Yeasen Biotechnology, #10651ES) was used to replace the unmodified UTP.

#### Circular mRNA production and purification

The *in vitro* transcription (IVT) of circular mRNA was carried out from linearized circular mRNA plasmid templates with the HiScribe T7 High Yield RNA Synthesis Kit (New England Biolabs, E2040S) following the kit manual. After IVT, circular mRNA was purified using the Monarch RNA Cleanup Kit (New England Biolabs, #T2050L). Then, the RNA precursors were heated to 70°C for 3 minutes and immediately placed on ice for 2 minutes. GTP was added to a final concentration of 2mM with a buffer (50mM Tris-HCl, 10mM MgCl2, 1mM DTT, pH 7.5) for 8 minutes at 55°C to catalyze the cyclization. Then RNA was column-purified.

For circular mRNA purification, we collected RNA fractions through UV absorbance at 260nm on an Agilent 1260 Series HPLC (Agilent) system with a 4.6×300mm column (Sepax Technologies, #215980P-4630) at a flow rate of 0.3mL/minute. The fractions were concentrated with a 4ml Ultracel-10 regenerated cellulose membrane (Millipore, #UFC8010) and purified by column chromatography. Then, the RNA was treated with RNase R (Beyotime, R7092L) for further enrichment. Finally, RNase R-digested RNA was column-purified.

### In vitro experiments

#### mRNA transfection

HEK293T cells were cultured in Dulbecco’s Modified Eagle’s Medium (DMEM) (BasalMedia, #L110KJ) containing 10% fetal bovine serum (NATOCOR, #SFBE) and 1% penicillin-streptomycin (GIBCO, #15140122) at the condition of 37°C and 5% CO_2_. mRNAs were transfected into HEK293T cells using Lipofectamine MessengerMax (Invitrogen, #LMRNA015) according to the manufacturer’s instructions. At the appropriate time after transfection, the cell lysate or supernatant was collected for protein detection.

#### *In vitro* protein expression measurements

For measurement of Gaussia luciferase activities, HEK293T cells were seeded in 96-well plate and transfected with 150ng mRNA per well. The cells were lysed at 6, 24, 48 and 72 hours after transfection using 1× cell lysis buffer from Dual Luciferase Reporter Assay Kit (Vazyme Biotech, DL101-01). Then, the luminescence signal was detected following the provided instructions.

For *in vitro* quantitative measurement of nerve growth factor (NGF), HEK293T cells were seeded in 24-well plate and 500ng of RNA was transfected into the cells per well. The cell culture supernatants were collected 6, 24, 48 and 72 hours after transfection. Then, the protein expression level was detected using Enzyme-linked Immunosorbent Assay Kit (Cloud-Clone Corp, #SEA105Mu), following the provided instructions.

For measurement of HA protein level, HEK293T cells were seeded in 12-well plate and transfected with 1.25ug mRNA per well. After 24 hours, cells were harvested and collected in 300ul RIPA lysis buffer (HANGZHOU DUDE BIOLOGICAL CO.LTD, #FD009) that contained 1% PMSF (HANGZHOU DUDE BIOLOGICAL CO.LTD, #FD0100). Then, protein level was analyzed with western blot using anti-flag primary antibody (Sigma-Aldrich, #F1804).

Each experiment was repeated three or four times from distinct samples.

#### Cell type specificity experiments

We used RiboDecode to design three Gluc mRNA variants optimized for preferential expression in HEK293T cells. The optimization process considered the cellular context of HEK293T cells while maintaining or reducing expression levels in A549 and ARPE19 cells. Gluc protein expression was measured 24 hours post-transfection. Each experiment was performed in quadruplicate. Expression levels were normalized to the reference (MF882921.1) Gluc mRNA for each cell type.

### In vivo experiments

#### Mouse retina histology and microscopy

For retinal whole-mounts immunofluorescence, eyes were surgically removed from perfused mice and fixed with 4% PFA at room temperature for 1 hour. Retinas were detached and whole mount staining was performed. The retinas were blocked for 1 hour in PBS staining buffer containing 5% normal donkey serum (Solarbio, SL050) and 0.1% Triton X-100 (Sigma, X100-100). The retinas were incubated with the primary antibody (Novus, NBP2-20112) overnight at 4°C and washed 3 times with PBS for 5 minutes each before incubation with the secondary antibody (CST, 4413S) for 2 hours at room temperature. The retinas were washed again with PBS 3 times for 5 minutes each and then mounted.

Confocal images were obtained using a Zeiss LSM 980 microscope. To count retinal ganglion cells (RGCs), we analyzed 320×320μm samples from the peripheral retina. These samples were taken ∼500μm from the center to the edge in all four quadrants of the retina. We then processed the data using ImageJ and ZEN software.

#### mRNA intravitreal injection

Adult mice were anesthetized by intraperitoneal injection of 1% sodium pentobarbital solution (25 mg/kg). Subsequently, a minor incision was made in the eyelid using a 30-gauge needle to facilitate eye exposure. For intravitreal injections, a micropipette was carefully inserted through the serosal opening, and formulations such as LNP-mRNA or other substances were administered into the vitreous body of the eye. To prevent the backflow of the injected solution, the needle was maintained in position for approximately 10 seconds after the injection before being gently withdrawn. To protect the cornea post-procedure, tobramycin was applied.

#### *In vivo* NGF protein expression

The m1Ψ-modified NGF mRNAs were injected into mouse retina and protein level were measured after 48 hours. The detachment and processing of the mouse retina were performed in the same way as mentioned in the section above. Then, the retinas were collected in 300ul RIPA lysis buffer (Beyotime, P0013B) that contained 1% PMSF (Sigma, 10837091001). Protein level was analyzed with western blot using anti-NGF Antibody-BSA and Azide free (Abcam, #ab6199).

#### Optic nerve crush mouse model

Mice were anesthetized by intraperitoneal injection of 1% sodium pentobarbital solution (25 mg/kg). Then, the eye surface was dilated with tropicamide drops and surface anesthesia was provided with proparacaine hydrochloride. The mice were fixed on the animal operating table. The optic nerve was completely exposed by cutting open the bulbar fascia under the surgical microscope and using microforceps to separate the surrounding tissues and hold the optic nerve for 5 s with a 0.07 mm wide reverse forceps at 1 mm posterior to the globe in the vertical direction of the longitudinal axis of the optic nerve. Tobramycin was applied daily to the superior orbital rim incision for 3 days postoperatively. Five to nine biological replicates were performed for each experiment.

#### *In vivo* immunogenicity

First, m^1^Ψ-modified mRNA with WT and RD1 codon sequences were encapsulated within lipid nanoparticles (LNP). BALB/c mice were received two intramuscular doses of 10μg mRNA each, with the dose determined by previous studies ^77,78^, administered on day 0 (prime) and day 14 (boost). (Figure 5d). We collected mouse serum and performed micro-neutralization (MN) assays to quantify neutralizing antibodies at two time points: day 14 (for prime response) and day 28 (for boost response) (Figure 5e, Methods). PBS buffer and the inactivated Split-virus (A/Victoria/2570/2019 (H1N1)) Influenza Vaccine was used as negative and positive control, respectively.

#### Micro-neutralization (MN) assay

To measure the titer of anti-influenza virus neutralizing antibodies, we treated mouse serum with receptor-destroying enzyme C (RDE C) (Denka-Seiken) at 37°C for 16 hours, followed by heat-inactivation at 56°C for 30 minutes. The experimental procedure of MN was the same as previously reported^79^. MN titration was defined as the reciprocal of the maximum serum dilution that neutralizes the 50% influenza H1N1/PR8 virus infections in MDCK cells. The minimum MN titer was set to 10. Eight biological replicates were performed for each experiment.

#### Ethics statement

For NGF mRNA *in vivo* assay, C57BL/6 mice were supplied by GemPharmatech. All experimental procedures involving these mice were conducted in strict accordance with the animal protocols that received approval from the Institutional Animal Care and Use Committee (IACUC) at the Zhongshan Ophthalmic Center, Sun Yat-Sen University, under the animal ethics approval number Z2021067. The *in vivo* experiments used male C57BL/6 mice aged 8 weeks. For HA mRNA *in vivo* assay, all the experimental procedures involving BALB/c mice were conducted in strict accordance with the animal protocols that received approval from the Animal Care and Use Committee Institutional at the Sun Yat-Sen University, under the animal ethics approval number 2022002794. The *in vivo* experiments used male BALB/c mice aged 6-8 weeks.

## DATA AVAILABILITY

The datasets used for modeling and analysis are listed in Supplementary file 1. The translation model and optimization framework of RiboDecode are available on GitHub (https://github.com/wangfanfff/RiboDecode) and figshare (https://figshare.com/articles/software/RiboDecode_zip/28916288?file=54138881). The codon sequences optimized by RiboDecode and UTR sequences in this study are listed in Supplementary file 2. Source Data are provided with this paper.

## ACKNOWLEDGMENTS

We gratefully acknowledge the researchers who made their Ribo-seq and RNA-seq data publicly available. We also extend our sincere appreciation to the reviewers for their insightful comments and suggestions, which have significantly improved this manuscript.

## AUTHORS’ CONTRIBUTIONS

Z.X. conceived, designed and supervised the project. YP.L. collected and preprocessed the datasets, evaluated the model performance and analyzed the data. Y. He, F.W., and ZH.T. developed the translation deep model. H.Z. developed the MFE deep model. F.W. and H.Z. developed the AM framework. JX.D. tested the model. JQ.Y. and LF.C. prepared mRNAs and conducted *in vitro* experiments. WB.J. conducted *in vivo* NGF experiments. ZR.H. and CJ.S. conducted the *in vivo* HA experiments. GF.Z encapsulated mRNA with LNP. X.H., H.J., Y. Hu, LP.W., and CY.Z. contributed to project discussion. YP.L., Z.X., Y. He and F.W. wrote the manuscript. All authors read and approved the manuscript.

