## Supplementary information for "Deep Generative Optimization of mRNA Codon Sequences for Enhanced mRNA Translation and Therapeutic Efficacy"

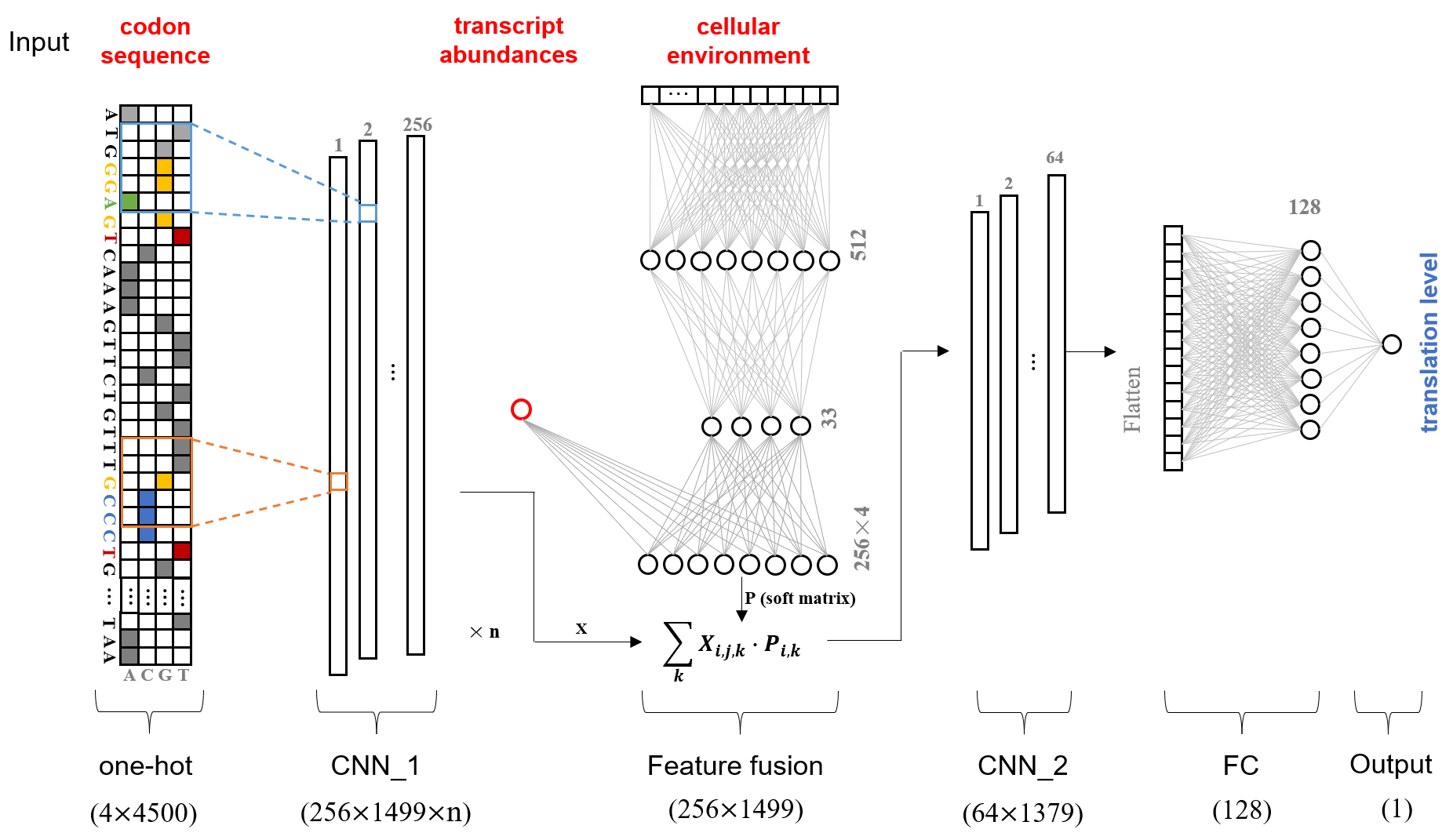

**Figure S1. Schematic diagram of the translation model structure.** The model accepts one-hot encoded nucleotide sequences as input, processed by convolutional neural networks (CNNs). A feature fusion mechanism is dynamically established through fully connected layers (FC) that integrate codon sequences, cellular environment features, and transcript abundance data (see Methods for implementation details). The dimensional shapes of each structural component are annotated in parentheses below the diagram. The number of CNN_1 was set to 4 for our model.

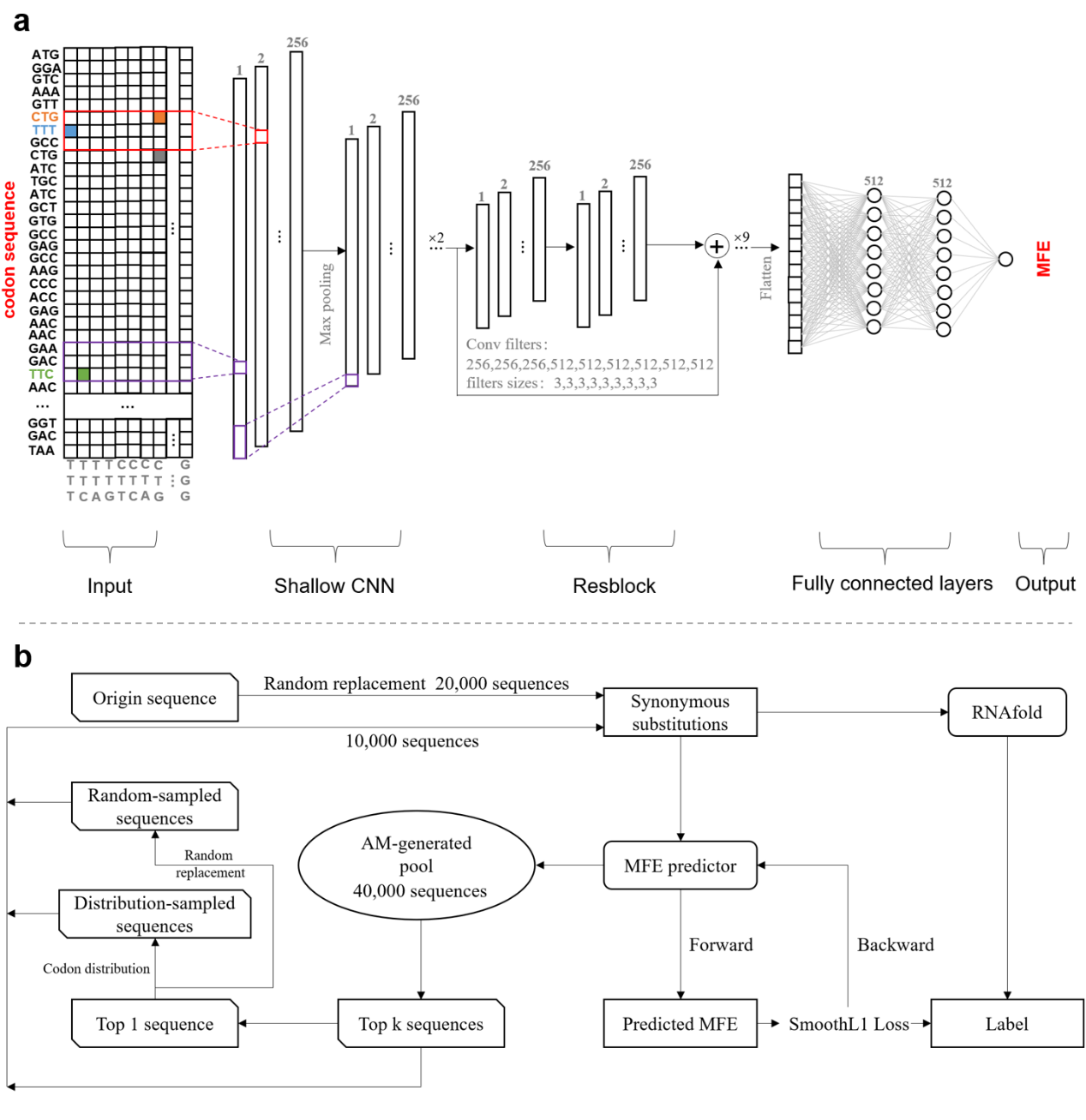

**Figure S2. MFE Model Architecture and Optimization Process**

**a.** MFE Model Architecture: a shallow CNN with two convolutional layers, nine Resblocks, and three fully connected layers (see Methods for details).

**b.** The MFE optimization contains four steps, including initial sampling, initial training, sequence generation and model retraining (see Methods for details).

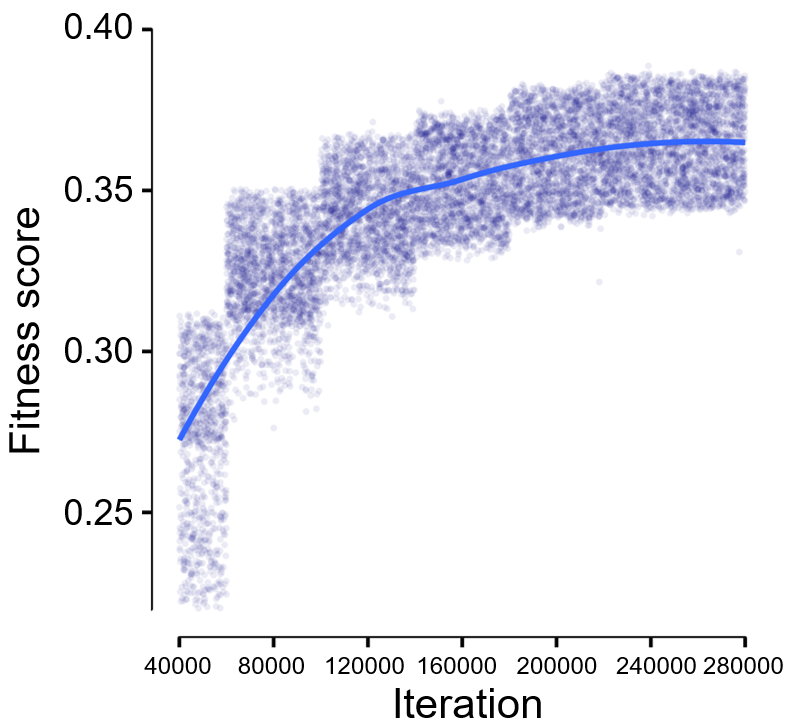

**Figure S3.** Fitness score trajectory during the iterative optimization of Gluc codon sequences (*w*=0.5). The fitness score is defined as the inverse of the loss function in the codon optimization framework (see Methods).

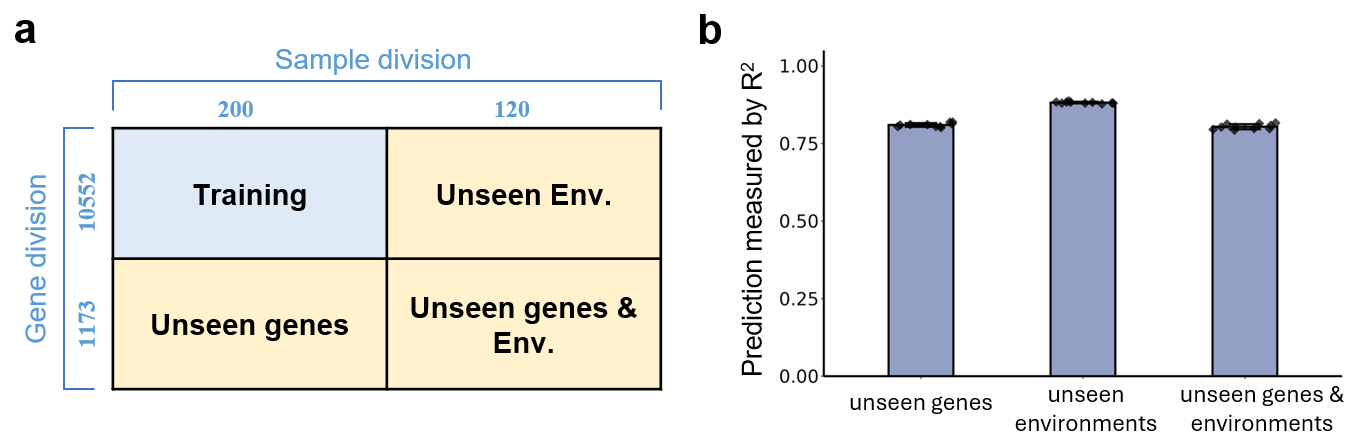

**Figure S4. Training and internal validation sets of the translation model.**

a. Schematic representation of how the dataset was divided into training (90% of genes) and test (10% of genes) sets. The 320 total datasets were split into 200 for training and 120 for validation, creating three validation sets: “unseen genes”, “unseen cellular environments”, and “unseen genes and cellular environments”.

b. 10-fold cross-validation on the test sets, where each dot on the bar represents a test. The error bars denote standard deviation.

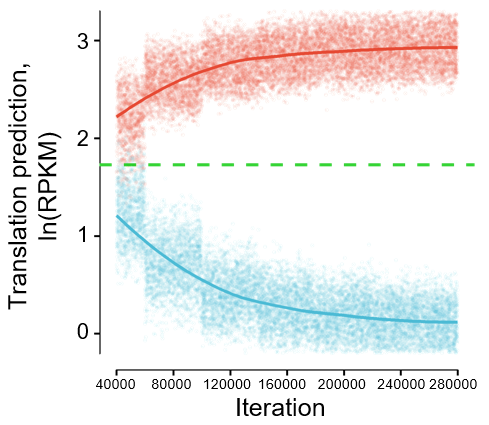

**Figure S5. Generated Gluc codon variants.** Plot showing the predicted translation levels of Gluc codon variants generated using the translation model (*w*=0). The x-axis represents generation iterations, with red and blue lines indicating enhanced and reduced predicted translation, respectively. The y-axis represents predicted translation level transformed to ln(RPKM$\times$5+1) (natural logarithm). The green dashed line shows the predicted translation level of the reference Gluc sequence.

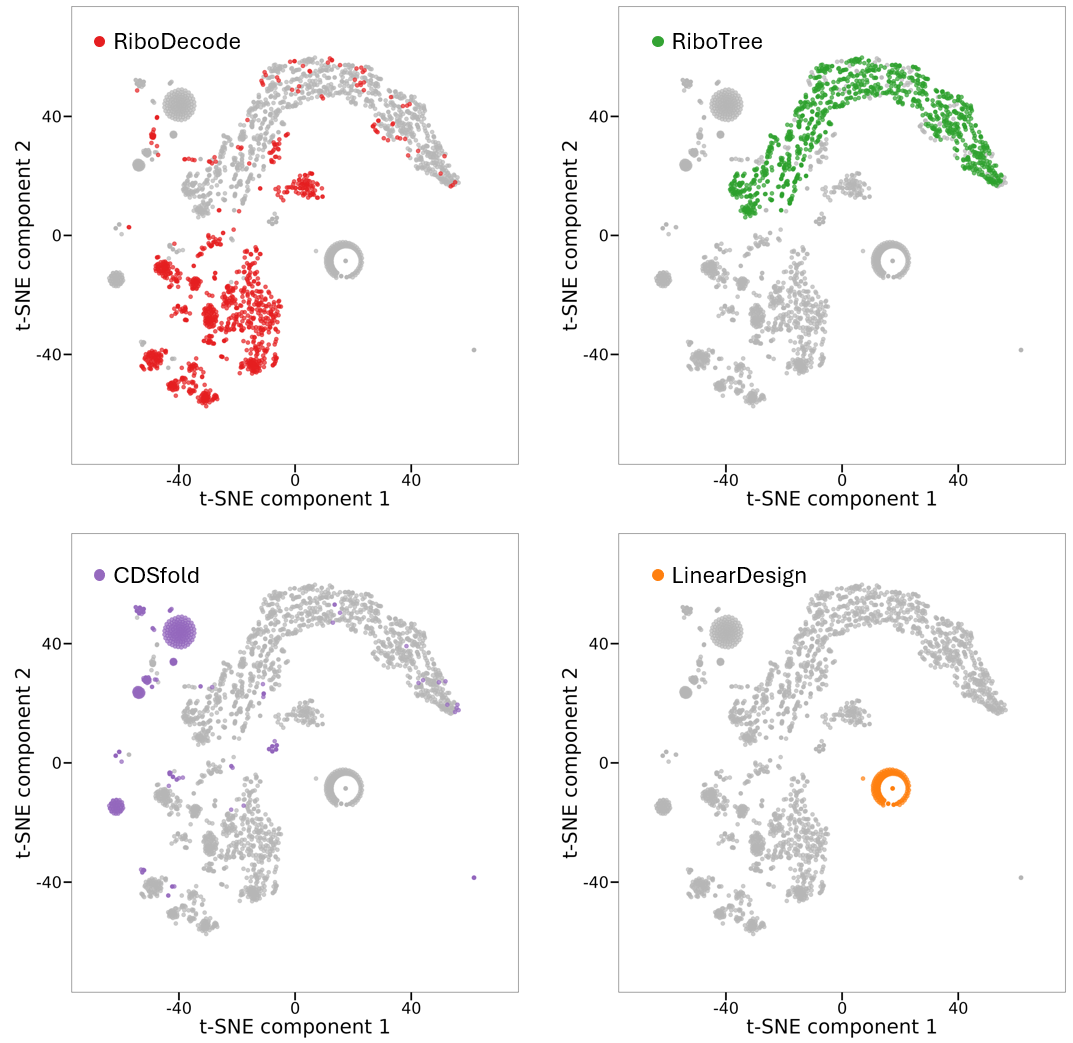

**Figure S6.** t-SNE distribution of Gluc codon sequences generated by RiboDecode, RiboTree, CDSfold, and LinearDesign (see Methods). The results show that the sequence space of RiboDecode is larger than other methods.

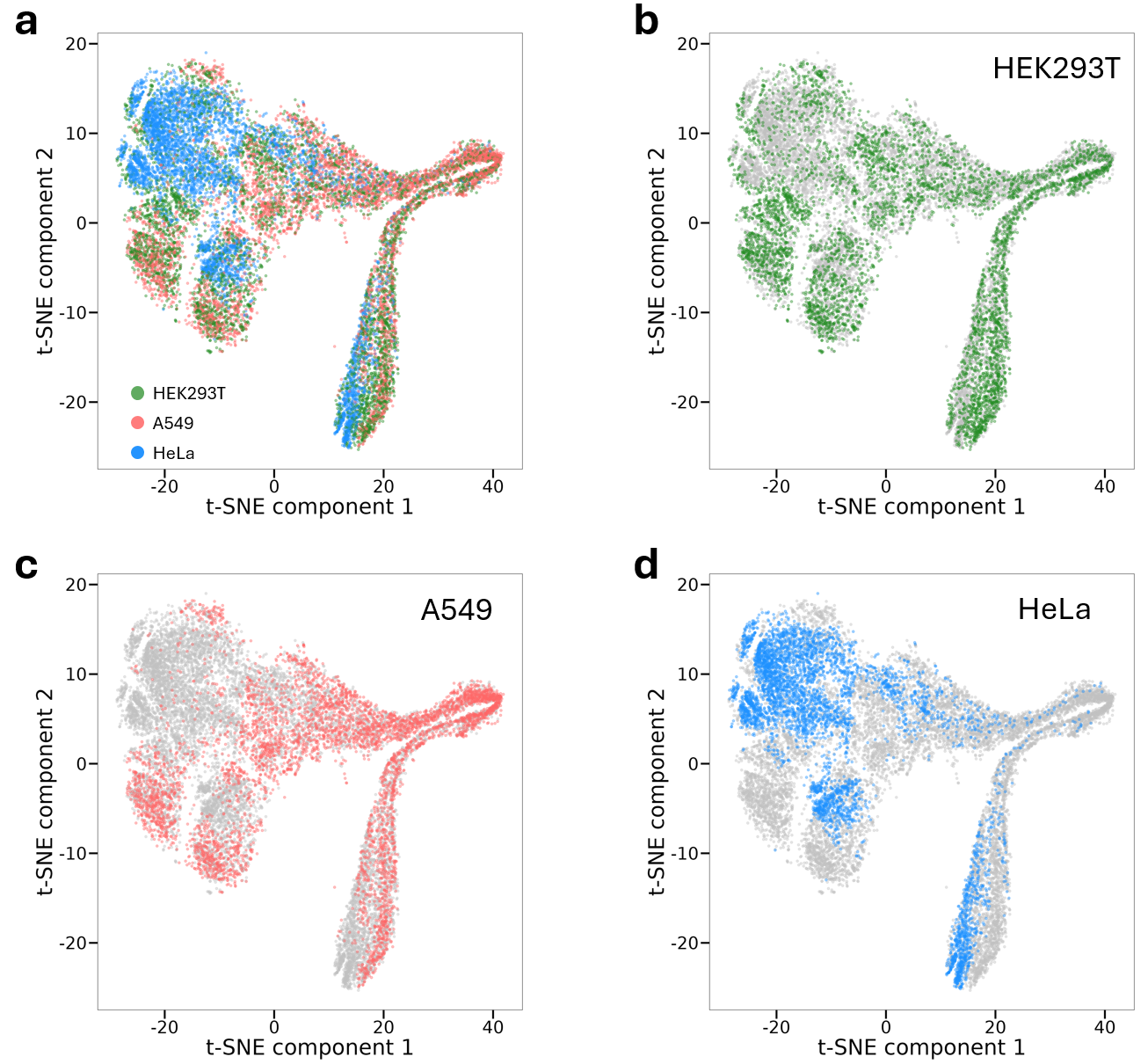

**Figure S7.** Gluc codon sequences were optimized by RiboDecode in different cellular contexts (*w*=0). The sequence embeddings were obtained using CodonBERT and visualized in a t-SNE plot. Each dot represents a single sequence, with the color indicating the cell type.

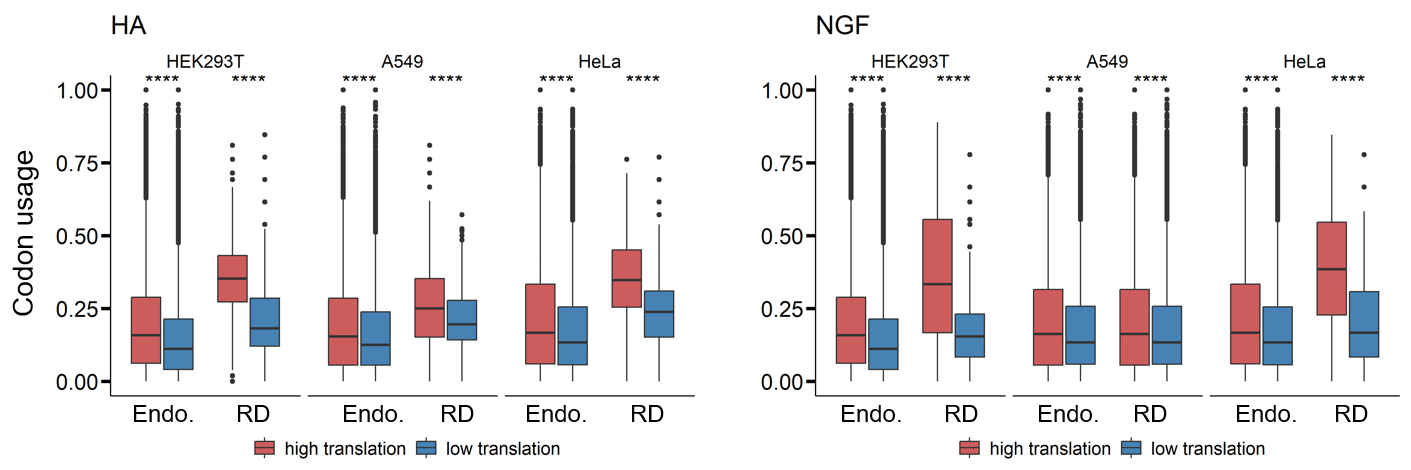

**Figure S8. Codon Usage Similarity Between Endogenous and RiboDecode-Generated Sequences.** Comparison of codon usage patterns in endogenous high-translation sequences and low-translation sequences, as well as the RiboDecode-designed sequences with enhanced and reduced translation for HA, and NGF in different cellular environments (HEK293T, A549, and HeLa). “Endo.”: endogenous. “RD”: RiboDecode-designed. (t-test, ****: *p*<0.0001).

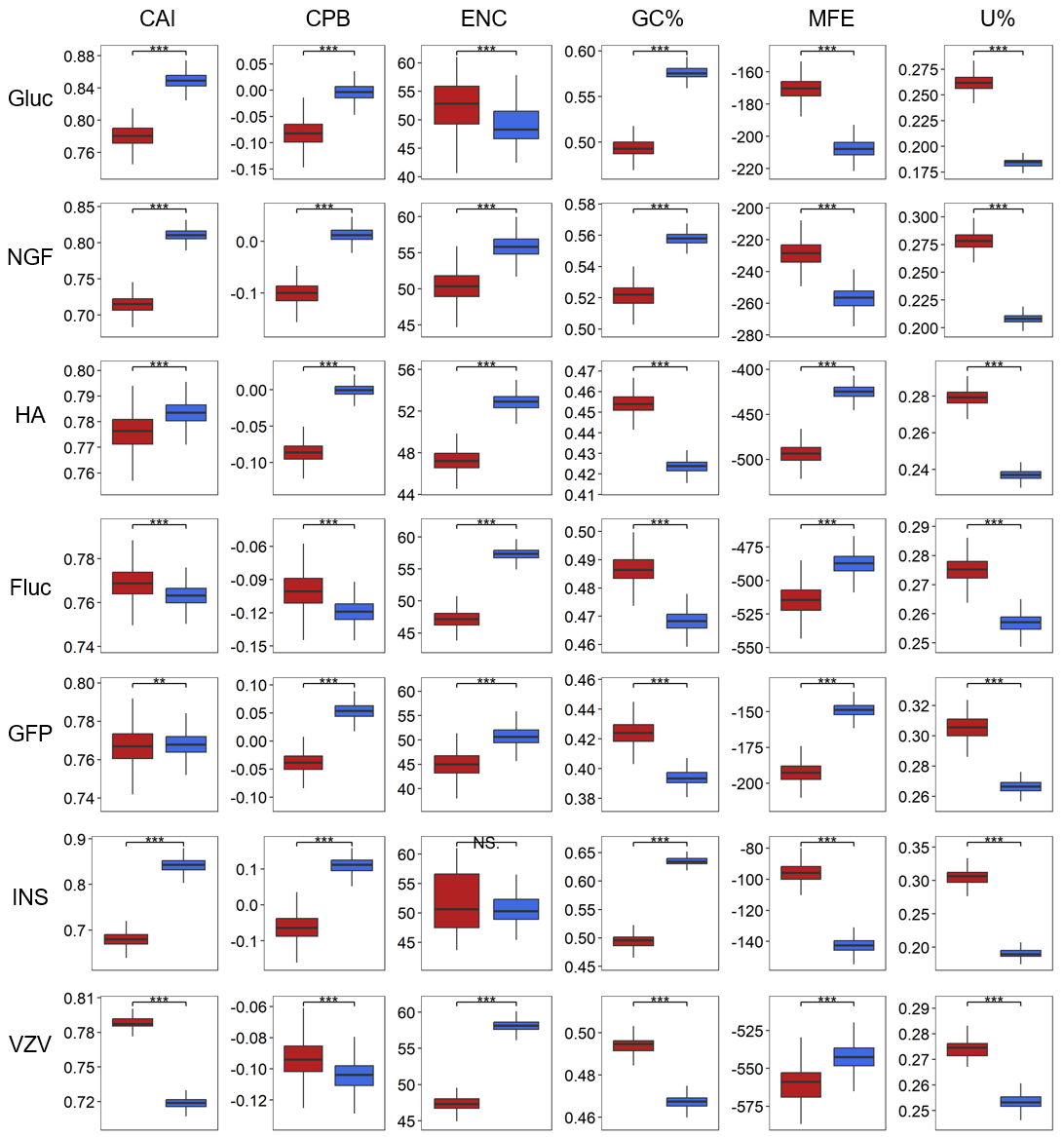

**Figure S9. Features of generated codon sequences for different genes.** The graphs show various sequence features for optimized (red) and unoptimized (blue) codon sequences for different genes (*w*=0). (t-test, ***: *p*<0.001). The full names of mRNAs are noted in the main text.

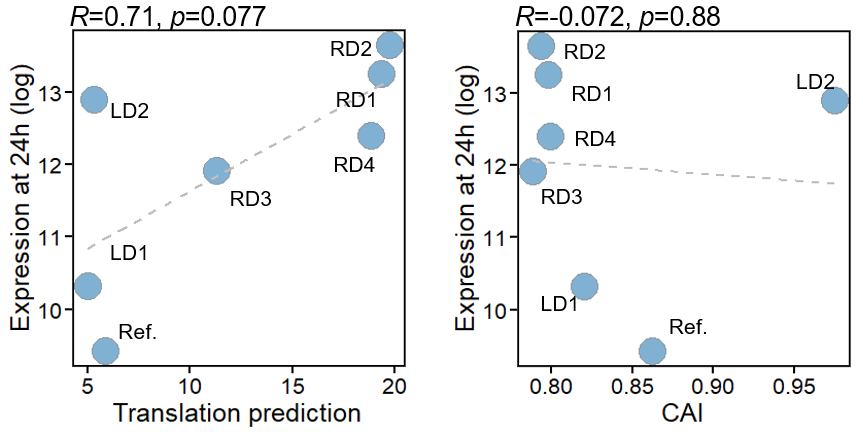

**Figure S10.** **Correlation of experimental expression with predicted measures.** The graphs show correlations between experimental protein expression (in natural logarithm) and translation predicted by RiboDecode, and CAI. Pearson’s correlation coefficients are provided.

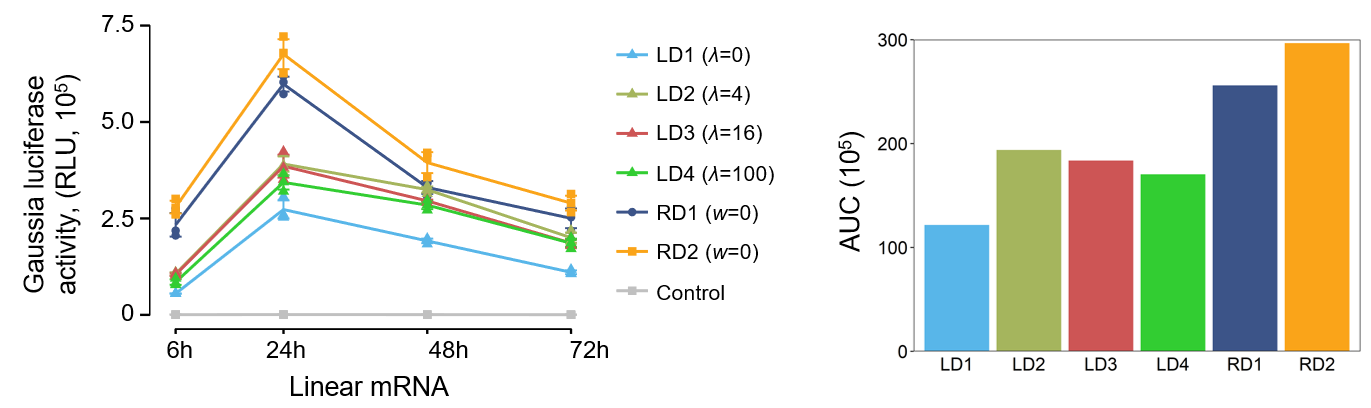

**Figure S11.** Protein expression of Gluc was measured by fluorescence intensity. RD1and RD2 were the same shown in Figure 4a, which had MFE values of -195.7 and -195.3 kcal/mol, respectively. LD1, LD2, LD3, and LD4 were designed using LinearDesign, with the MFEs of -346.2, -302.5, -266.8, and -246.7 kcal/mol, respectively. “RLU”: relative light units. The area under the curve (AUCs) were calculated.

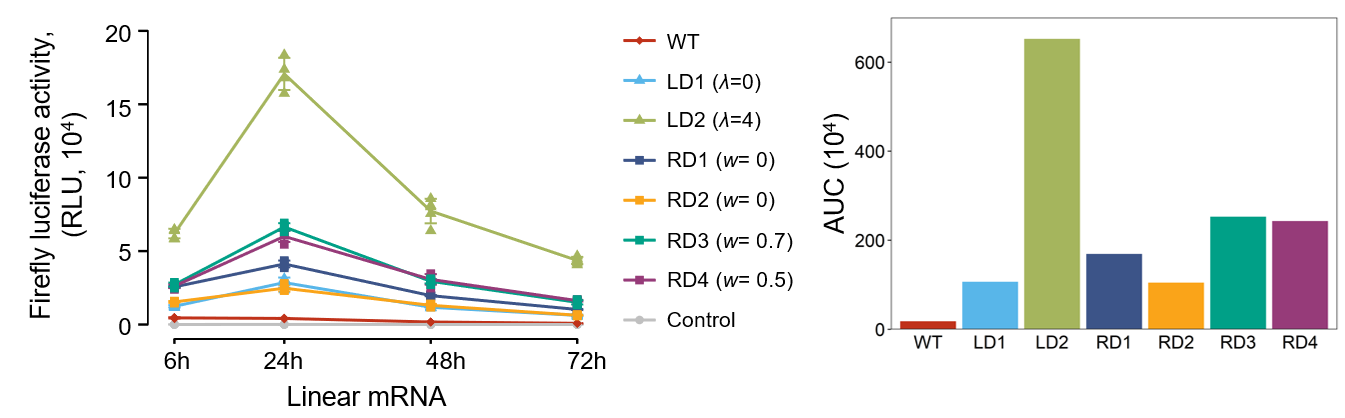

**Figure S12. Fluc protein expression in transfected HEK293T cells.** Protein expression was determined by fluorescence intensity. RD1, RD2, RD3, and RD4 were designed using RiboDecode, with the *w* parameters indicated in brackets. The detailed information of the sequences is provided in Table S6. “RLU”: relative light unit. The area under the curve (AUCs) were calculated.

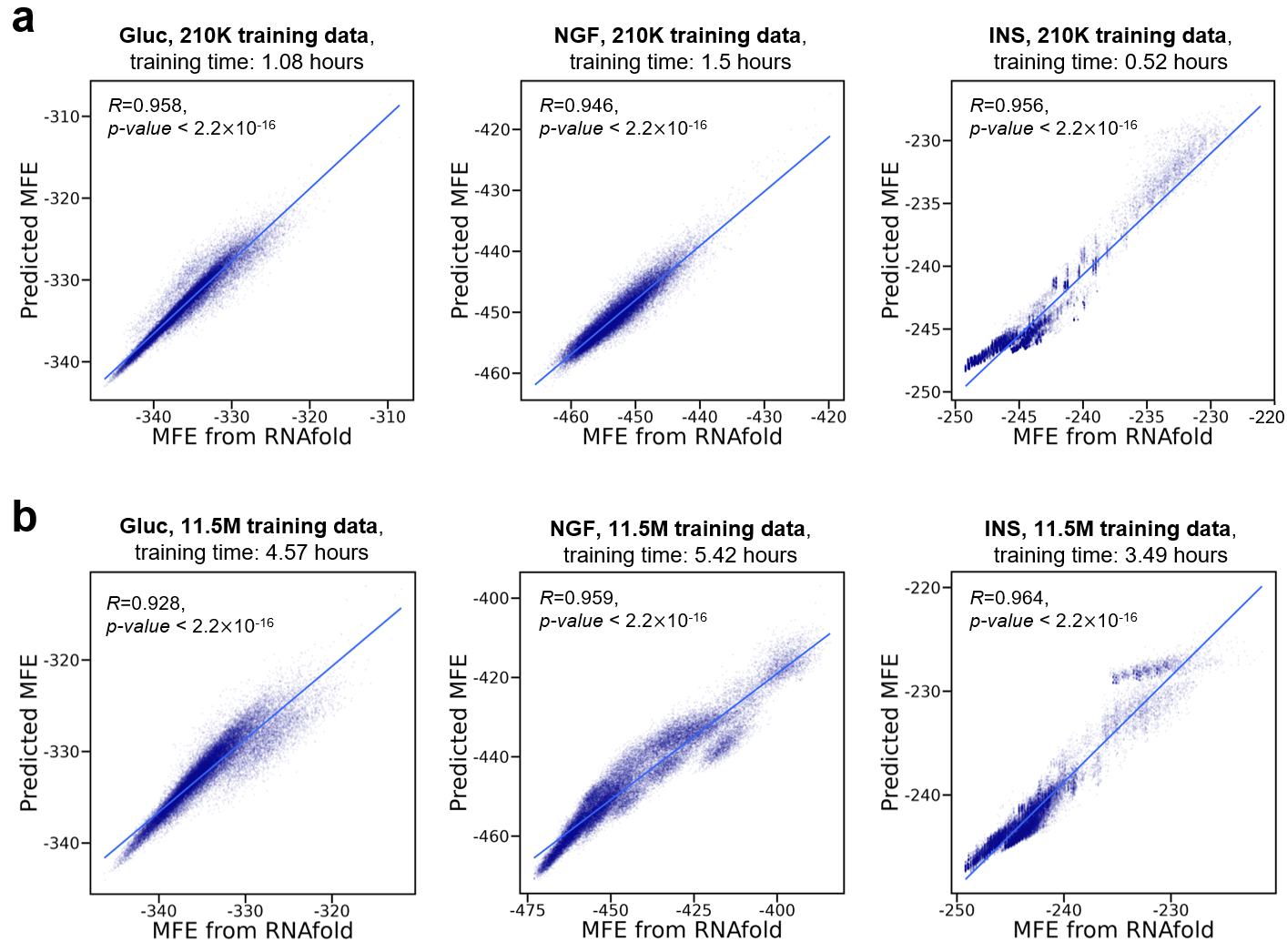

**Figure S13. Evaluation of the Predictive Performance of the MFE Model to RNAfold.** a. MFE models for different mRNAs trained by 210K samples. b. MFE models for different mRNAs trained by 11.5M samples.

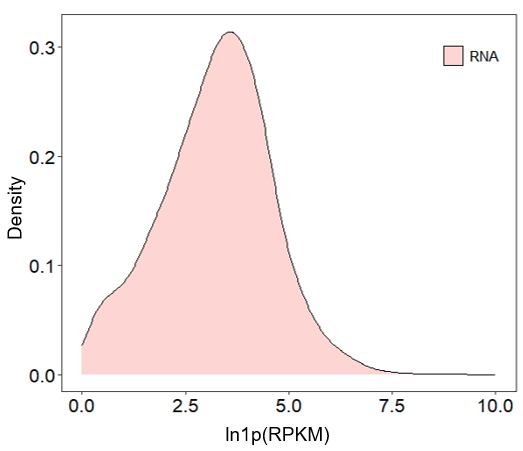

**Figure S14. Distribution of mRNA Abundance Levels Used for Modeling.** The graph shows the distribution of mRNA expression for 11,725 genes in 320 samples. Expression counts were transformed to ln(y×RPKM+1), where y was set to 5 to maximize the correlation between mRNA abundance and translation level. RPKM stands for reads per kilobase of transcript per million reads mapped. The default mRNA count for codon optimization was set to 4.5 based on the median value of this distribution.

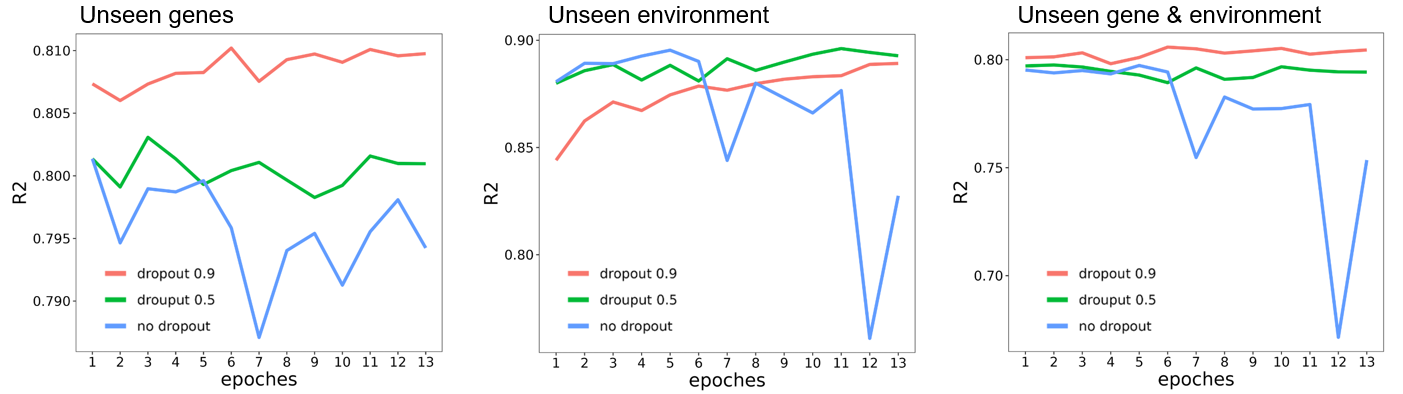

**Figure S15. Evaluation of translation prediction models trained with varying dropout rates. R² values across three test sets are shown.**

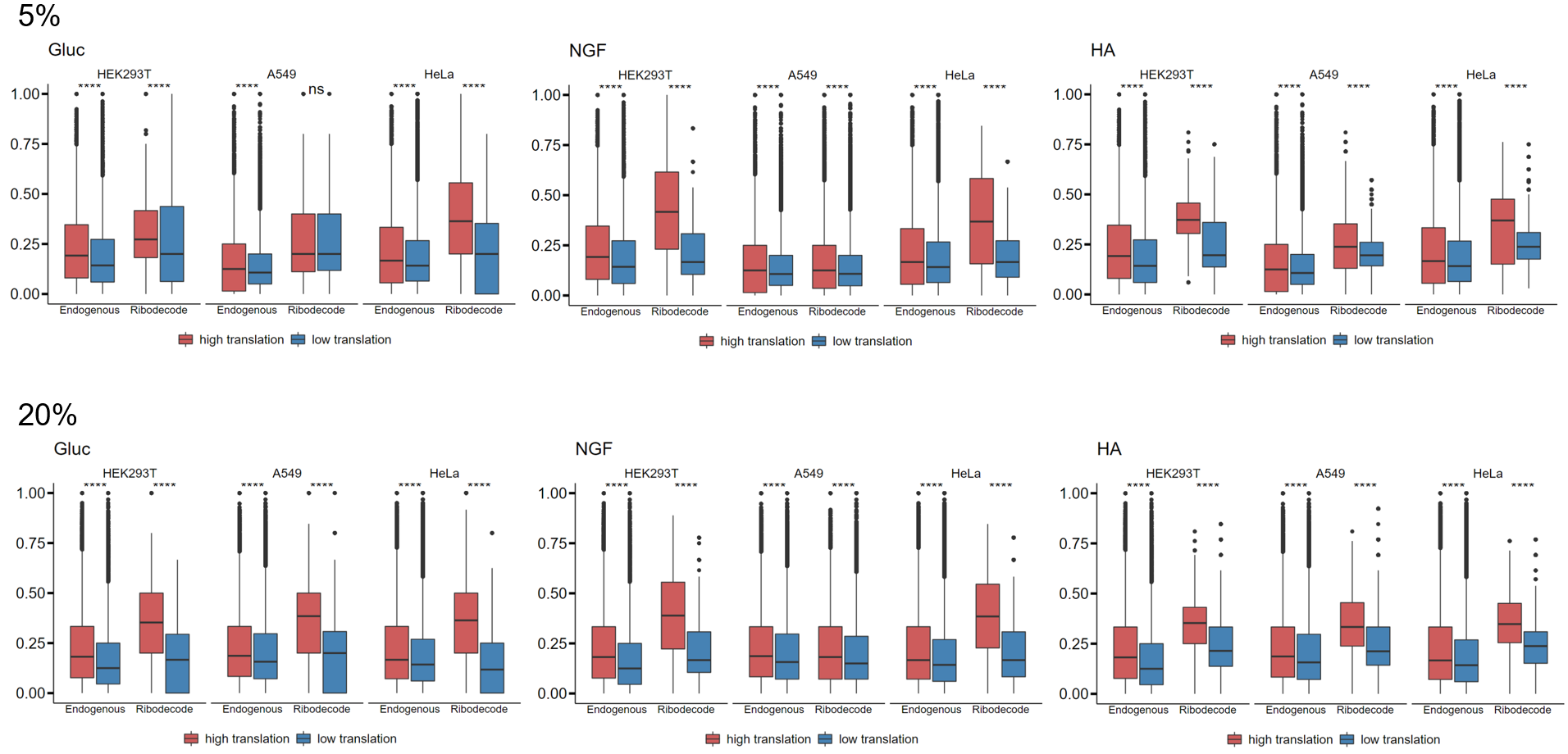

**Figure S16. Codon Usage Similarity Between Endogenous and RiboDecode-Generated Sequences in thresholds of 5% and 20%.** Comparison of codon usage patterns in endogenous high-translation sequences and low-translation sequences, as well as the RiboDecode-designed sequences with enhanced and reduced translation for HA, and NGF in different cellular environments (HEK293T, A549, and HeLa). “Endo.”: endogenous. “RD”: RiboDecode-designed. (t-test, ****: *p*<0.0001).

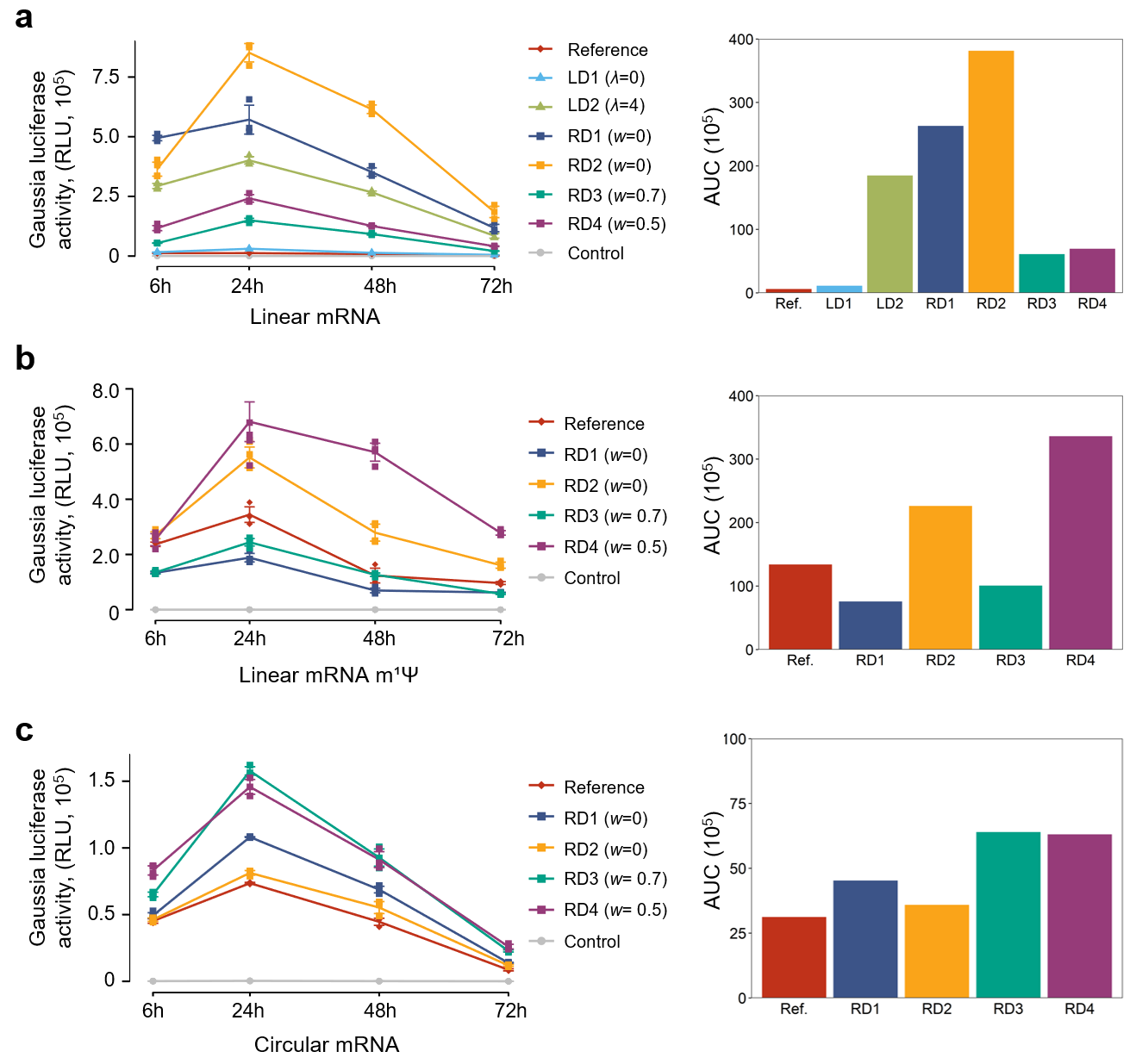

**Figure S17.** Protein expression of Gluc. The area under the curve (AUCs) were calculated based on Figure 4.

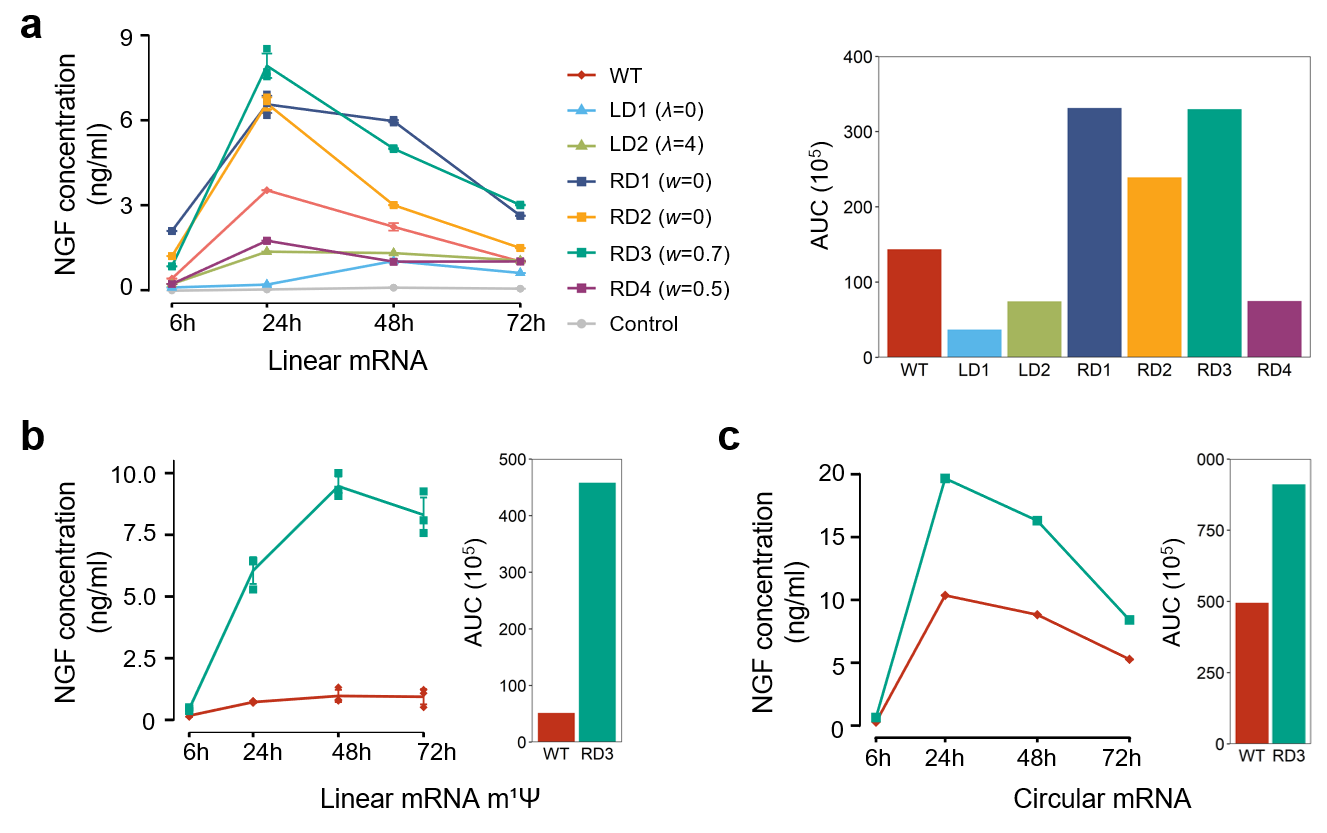

**Figure S18.** Protein expression of NGF. The area under the curve (AUCs) were calculated based on Figure 6.

| **Input** | **Training set (R^2^)** | **Test set (R^2^)** | | |
| --- | --- | --- | --- | --- |
|  |  | **New gene** | **New Env.** | **New gene & Env.** |
| **basic** | **0.86021** | **0.81329** | **0.88719** | **0.80838** |
| + codon frequency | 0.86108 | 0.800528 | 0.883919 | 0.790282 |
| + MFE | 0.86086 | 0.809827 | 0.889674 | 0.800357 |
| + CAI | 0.86127 | 0.802417 | 0.880232 | 0.791394 |
| + cell type | 0.8609 | 0.808396 | 0.890336 | 0.799348 |
| + treatment | 0.86174 | 0.803956 | 0.885125 | 0.796533 |

**Table S1. Evaluation of translation prediction model on internal test sets.** This table compares the performance (R²) of the basic translation model with models incorporating additional inputs on various test sets. The basic input includes codon sequence, mRNA abundance, and cellular environment represented by gene expression profiles from RNA-seq. “Env.” stands for cellular environment.

| **Input** | **Training set (R^2^)** | **Test set (R^2^)** |
| --- | --- | --- |
| CDS + mRNA abundances + cellular environment | 0.85924 | 0.819247 |
| mRNA abundances + cellular environment | 0.7331 | 0.62351 |
| CDS + mRNA abundances | 0.80825 | 0.760404 |
| CDS + cellular environment | 0.58303 | 0.467216 |
| CDS | 0.56135 | 0.429161 |
| mRNA abundances | 0.70318 | 0.608484 |

**Table S2. Ablation analysis of translation model inputs.** This table shows the results of an ablation analysis investigating the contribution of the three main inputs (codon sequences, mRNA abundances, cellular environment) to the translation model’s performance. “CDS”: codon sequence.

| **Gene** | **Sequence ID** | **Parameter** | **Predicted translation level** | **MFE** | **CAI** |
| --- | --- | --- | --- | --- | --- |
| Gluc | Reference | NA | 5.88 | -216.4 | 0.862 |
| Gluc | RD1 | w=0 | 19.34 | -195.7 | 0.798 |
| Gluc | RD2 | w=0 | 19.75 | -195.3 | 0.794 |
| Gluc | RD3 | w=0.7 | 11.28 | -342.4 | 0.788 |
| Gluc | RD4 | w=0.5 | 18.85 | -258.4 | 0.799 |
| Gluc | LD1 | λ=0 | 4.99 | -346.2 | 0.821 |
| Gluc | LD2 | λ=4 | 5.29 | -302.5 | 0.975 |
| Gluc | LD3 | λ=16 | 4.33 | -266.8 | 0.991 |
| Gluc | LD4 | λ=100 | 4.40 | -246.7 | 0.993 |

**Table S3.** **Sequence information of Gluc mRNA variants.** This table presents data for various Gluc mRNA constructs, including the reference sequence, RiboDecode-generated variants (RD1-RD4 with different *w* values), and LinearDesign-generated variants (LD1-LD4). All MFE values are reported in kcal/mol.

| **Sequence ID** | **Protein expression level (24h)** | | | **Relative expression (prediction; experiments)** | |
| --- | --- | --- | --- | --- | --- |
|  | **In HEK293T** | **In A549** | | **In HEK293T** | **In A549** |
| Gluc-Reference | 24527.5 | | 28271.25 | 1; 1 | 1; 1 |
| HEK_A549_1 | 38522.25 | | 32671 | 2.16; 1.57 | 1.25; 1.16 |
| HEK_A549_2 | 29584 | | 19712.75 | 1.98; 1.21 | 1.10; 0.70 |
| HEK_A549_3 | 23215.25 | | 21544 | 2.32; 1.29 | 1.41; 0.76 |

**Table S4.** **Experimental protein expression of Gluc mRNA variants with cellular differential expression.** This table presents experimental protein expression for various Gluc mRNA construct, including the reference sequence and RiboDecode-designed variants. The designed variants were predicted to express more in HEK293T than in A549 (HEK_A549_1 to 3, *w*=0). For each construct, the table shows original protein expression level, predicted expression relative to the reference, and protein expression level relative to the reference. Protein expression was measured by fluorescence intensity.

| **Sequence ID** | **Protein expression level (24h)** | | **Relative expression (prediction; experiments)** | |
| --- | --- | --- | --- | --- |
|  | **In HEK293T** | **In ARPE19** | **In HEK293T** | **In ARPE19** |
| Gluc-Reference | 24527.5 | 106308.5 | 1; 1 | 1; 1 |
| HEK_ARPE_1 | 23215.25 | 34516 | 2.02; 0.95 | 1.43; 0.32 |
| HEK_ARPE_2 | 17301 | 23928.5 | 2.23; 0.71 | 1.59; 0.23 |
| HEK_ARPE_3 | 31671.25 | 53428.25 | 1.42; 1.29 | 0.99; 0.50 |

**Table S5.** **Experimental protein expression of Gluc mRNA variants with cellular differential expression.** This table presents experimental protein expression for various Gluc mRNA construct, including the reference sequence and RiboDecode-designed variants. The designed variants were predicted to express more in HEK293T than in ARPE19 (HEK_ARPE_1 to 3, *w*=0). For each construct, the table shows original protein expression level, predicted expression relative to the reference, and protein expression level relative to the reference. Protein expression was measured by fluorescence intensity.

| **Gene** | **Sequence ID** | **Parameter** | **Predicted translation level** | **MFE** | **CAI** |
| --- | --- | --- | --- | --- | --- |
| Fluc | Reference | NA | 7.20 | -216.4 | 0.808 |
| Fluc | RD1 | w = 0 | 12.94 | -195.7 | 0.713 |
| Fluc | RD2 | w = 0 | 12.01 | -195.3 | 0.703 |
| Fluc | RD3 | w = 0.7 | 9.34 | -342.4 | 0.748 |
| Fluc | RD4 | w = 0.5 | 11.47 | -258.4 | 0.736 |
| Fluc | LD1 | λ=0 | 4.45 | -346.2 | 0.766 |
| Fluc | LD2 | λ=4 | 5.03 | -302.5 | 0.952 |

**Table S6.** This table presents sequence features for various Fluc mRNA constructs, including the wild-type sequence, RiboDecode-generated variants (RD1-RD4 with different *w* values), and LinearDesign-generated variants (LD1-LD2). All MFE values are reported in kcal/mol.

| **Gene** | **Sequence ID** | **Parameter** | **Predicted translation level** | **MFE** | **CAI** |
| --- | --- | --- | --- | --- | --- |
| HA | WT | NA | 3.39 | -411.8 | 0.788 |
| HA | RD1 | w = 0 | 28.93 | -535.7 | 0.773 |
| HA | RD2 | w = 0 | 29.76 | -536.8 | 0.783 |
| HA | RD3 | w = 0.7 | 10.41 | -964.4 | 0.794 |
| HA | RD4 | w = 0.5 | 12.53 | -885.7 | 0.782 |
| HA | LD1 | λ=0 | 3.58 | -1098.6 | 0.773 |
| HA | LD2 | λ=4 | 3.19 | -899.5 | 0.961 |

**Table S7.** Predicted translation levels and experimental protein expression of HA mRNA variants. This table presents data for various HA mRNA constructs, including the wild-type sequence, RiboDecode-generated variants (RD1-RD4 with different *w* values), and LinearDesign-generated variants (LD1-LD2). All MFE values are reported in kcal/mol.

| **Gene** | **Sequence ID** | **Parameter** | **Predicted translation level** | **MFE** | **CAI** |
| --- | --- | --- | --- | --- | --- |
| NGF | WT | NA | 5.08 | -262.5 | 0.821 |
| NGF | RD1 | w = 0 | 17.52 | -251.6 | 0.761 |
| NGF | RD2 | w = 0 | 17.14 | -250.3 | 0.755 |
| NGF | RD3 | w = 0.7 | 10.89 | -432.3 | 0.728 |
| NGF | RD4 | w = 0.5 | 14.94 | -366 | 0.737 |
| NGF | LD1 | λ=0 | 3.96 | -523.8 | 0.741 |
| NGF | LD2 | λ=4 | 3.19 | -442.5 | 0.953 |

**Table S8.** Predicted translation levels and experimental protein expression of NGF mRNA variants. This table presents data for various NGF mRNA constructs, including the wild-type sequence, RiboDecode-generated variants (RD1-RD4 with different *w* values), and LinearDesign-generated variants (LD1-LD2). All MFE values are reported in kcal/mol.

| **Category** | **Parameter Name** | **Description** | **Values** | **Value Range** |
| --- | --- | --- | --- | --- |
| Training Configuration | num_epochs | Number of training epochs | 20 | fixed |
|  | learning_rate | Initial learning rate | 0.001 | fixed |
|  | lr_scheduler_milestones | Epochs when learning rate will drop | [10] | fixed |
|  | lr_scheduler_gamma | Learning rate decay factor at each milestone | 0.4 | fixed |
|  | weight_decay | L2 regularization coefficient | 0.01 | fixed |
|  | bias_initialization | Custom bias parameter initialization | (0, 1) | random variable |
|  | batch_size | Training batch size | 100 | fixed |
| Data & parameter | training_data | Number of training data | 2.11M | fixed |
|  | model_parameters | Total trainable parameters | 17.27M | fixed |
| CNN Architecture | conv1d_kernel_size | 1D convolution kernel size | [5, 30] | fixed |
|  | conv1d_filters | Number of 1D convolution filters | [64, 256] | fixed |
|  | conv1d_dilation | Dilation rate for 1D convolution | [1, 4] | fixed |
|  | conv1d_stride | Stride for 1D convolution | [1, 3] | fixed |
| Pooling & Regularization | maxpooling_kernel_size | Max-pooling kernel size | 3 | fixed |
|  | maxpooling_stride | Max-pooling stride | 1 | fixed |
|  | dropout_rate | Dropout probability | [0.5, 0.9] | fixed |
| Fully Connected | output_layers | Number of final output layers | 2 | fixed |
|  | dense_layer_units | Fully connected layer dimensions | [32, 128, 512] | fixed |
|  | dropout_rate | Dropout probability | 0.025 | fixed |

**Table S9.** **Hyperparameters for the translation model.** This table lists the hyperparameters used in the translation prediction model. It includes global parameters and specific parameters for the Convolutional Neural Network and Fully Connected layers. The table specifies whether each parameter is fixed or variable within a given range.

| **Category** | **Parameter Name** | **Description** | **Values** | **Value Range** |
| --- | --- | --- | --- | --- |
| Training Configuration | num_epochs | Number of training epochs | 20 | fixed |
|  | learning_rate | Initial learning rate | 0.001 | fixed |
|  | lr_scheduler_milestones | Epochs when learning rate will drop | [1] | fixed |
|  | lr_scheduler_gamma | Learning rate decay factor at each milestone | 0.1 | fixed |
|  | weight_decay | L2 regularization coefficient | 5e-4 | fixed |
|  | epsilon_greedy | Epsilon parameter for exploration | (0.1, 1) | random variable |
|  | batch_size | Training batch size | 128 | fixed |
| Data & parameter | training_data | Number of training data | 210,000 | fixed |
|  | model_parameters | Total trainable parameters | 10.88M | fixed |
| CNN Architecture | conv1d_kernel_size | 1D convolution kernel size | [1, 3] | fixed |
|  | conv1d_filters | Number of 1D convolution filters | [128, 256] | fixed |
|  | conv1d_dilation | Dilation rate for 1D convolution | 1 | fixed |
|  | conv1d_stride | Stride for 1D convolution | 1 | fixed |
| Pooling & Regularization | maxpooling_kernel_size | Max-pooling kernel size | 3 | fixed |
|  | maxpooling_stride | Max-pooling stride | 2 | fixed |
|  | dropout_rate | Dropout probability | 0.5 | fixed |
|  | leaky_relu_slope | Negative slope for LeakyReLU | 0.2 | fixed |
| ResNet Components | residual_blocks | Number of BasicBlock residual units | 8 | fixed |
|  | adaptive_pooling_output | Adaptive average pooling output size | 1 | fixed |
| Fully Connected | output_layers | Number of final output layers | 2 | fixed |
|  | dense_layer_units | Fully connected layer dimensions | [256, 512] | fixed |

**Table S10.** **Hyperparameters for MFE model.** This table presents the hyperparameters used in the MFE prediction model. Like Table S9, it includes global parameters and specific parameters for the CNN and Fully Connected layers. The table indicates whether each parameter is fixed or variable within a specified range, providing insights into the model’s architecture and training process.

| **Type** | **Parameter name** | **Values** | **Value range** |
| --- | --- | --- | --- |
| Global | α | 100 | fixed |
|  | β | 100 | fixed |
|  | parameters | 274.30 K | fixed |

**Table S11.** Hyperparameters for Codon Optimizer.

| **Gene** | **Items** | **JAX-RNAfold** | | **Our MFE model** |
| --- | --- | --- | --- | --- |
| Gluc (558 nt) | Time-cost | 1.55 hours | 1.08 hours | |
|  | GPU memory | 26.36 GB | 1,012 MB | |
|  | MFE | -327.1 kcal/mol | -345.46 kcal/mol | |

**Table S12.** Gluc MFE optimization by JAX-RNAfold and our model. The MFE values were calculated by RNAfold.
